# A complete allosteric map of a GTPase switch in its native network

**DOI:** 10.1101/2022.04.13.488230

**Authors:** Christopher J.P. Mathy, Parul Mishra, Julia M. Flynn, Tina Perica, David Mavor, Daniel N.A. Bolon, Tanja Kortemme

## Abstract

Allosteric regulation is central to protein function in cellular networks^1^. However, despite technological advances^2,3^ most studies of allosteric effects on function are conducted in heterologous environments^2,4,5^, limiting the discovery of allosteric mechanisms that rely on endogenous binding partners or posttranslational modifications to modulate activity. Here we report an approach that enables probing of new sites of allosteric regulation at residue-level resolution in essential eukaryotic proteins in their native biological context by comprehensive mutational scanning. We apply our approach to the central GTPase Gsp1/Ran. GTPases are highly regulated molecular switches that control signaling, with switching occurring via catalyzed GTP hydrolysis and nucleotide exchange. We find that 28% of 4,315 assayed mutations in Gsp1/Ran are highly deleterious, showing a toxic response identified by our assay as gain-of-function (GOF). Remarkably, a third of all positions enriched for GOF mutations (20/60) are outside the GTPase active site. Kinetic analysis shows that these distal sites are allosterically coupled to the active site, including a novel cluster of sites that alter the nucleotide preference of Gsp1 from GDP to GTP. We describe multiple distinct mechanisms by which allosteric mutations alter Gsp1/Ran cellular function by modulating GTPase switching. Our systematic discovery of new regulatory sites provides a functional map relevant to other GTPases such as Ras that could be exploited for targeting and reprogramming critical biological processes.

## Main Text

Allostery, the process by which perturbations at one site of a protein exert functional effects at distal sites, is a central regulatory mechanism in cells^1^. Protein or ligand binding, posttranslational modifications, and mutations can allosterically alter subsequent binding events or enzymatic activities to control metabolism^3^ or signaling^6,7^, making allosteric regulation a driver of disease and attractive target for therapeutic drug design^8^. While it has been suggested that a considerable fraction of protein residues may be primed for allosteric regulation^2^ and this priming may enable the evolution of new functional protein-protein interactions^9^, it remains an open question how prevalent allosteric sites are in a protein structure. Moreover, while biophysical aspects of allostery have been mapped using technological advances^2^, the role of allosteric perturbations on cellular function in physiological networks has not been mapped comprehensively even for single proteins. One contributor is a lack of methods for discovering new sites of allosteric regulation in the cellular context, thus limiting the identification of new targets for drug development and the reprogramming of functions in cellular networks.

A class of proteins thought to be regulated through allosteric mechanisms are switches, which cycle between “on” and “off” states in response to signals, are ubiquitous in biological regulation^10^, and whose misregulation is often associated with disease^11^. In small GTPase switches, interconversion between a GTP-bound on-state and a GDP-bound off-state is intrinsically slow but is accelerated by two opposing regulators: GTPase-activating (GAP) proteins that activate GTP hydrolysis and guanine nucleotide exchange factor (GEF) proteins that accelerate nucleotide replacement. Perturbations at a very limited number of allosteric sites distal from the active site, which comprises the nucleotide binding region and the switch loops^12^, have been shown to affect the kinetics of biochemical switching function *in vitro^6^* and to lead to switch overactivation^4,5^ and altered cellular function^*6*^. Additionally, one allosteric site of the GTPase Ras has been successfully targeted by small molecule inhibitors^13^. Despite these key findings, the vast majority of GTPase sites remain untested for allosteric regulation in their native biological networks^14^ when the functional context of opposing regulators, posttranslational modifications, interaction partners, and downstream signaling pathways is preserved (**Fig. 1a**).

**Figure 1.**
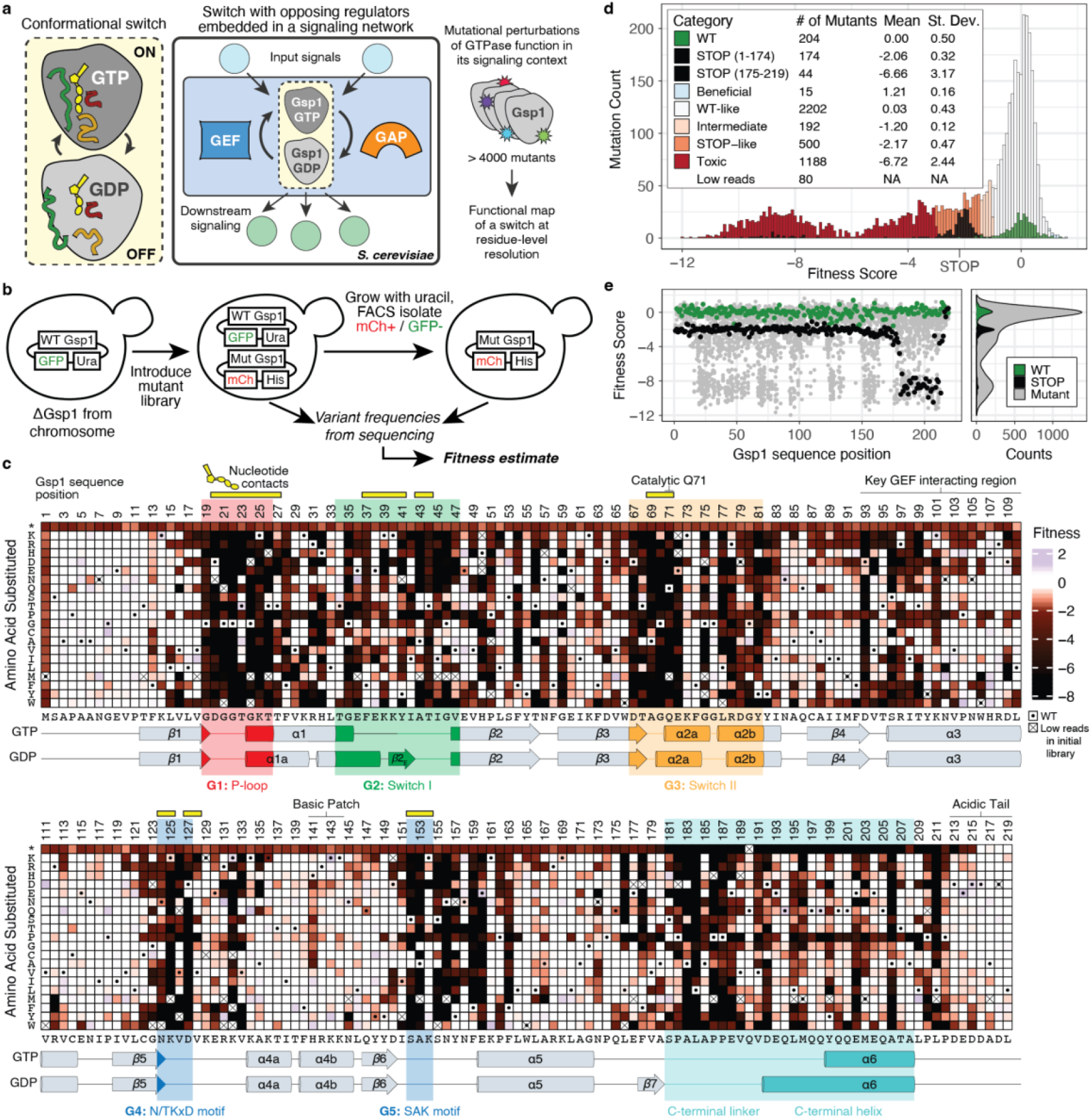
*In vivo* sensitivity of the GTPase Gsp1 to all possible single amino acid substitutions. **a,** Mutational perturbations exhaustively probe a switch in its native network. **b,** Generalizable plasmid swap approach to probe essential genes by mutational mapping. **c,** Heatmap showing quantitative fitness scores for all Gsp1 mutations after 6 generations of competitive growth. Dot indicates WT synonymous codons; X indicates mutants with low reads in the initial library outgrowth. Conserved G1-5 regions are shown in colors corresponding to structural annotations in **Fig. S1**. Additional annotated functional regions include the catalytic residue Q71, the GEF interacting region, and the basic patch and acidic tail that interact in the GDP-bound structure^24^. Positions contacting the nucleotide or magnesium cofactor are indicated by yellow bars. Secondary structure assignments for each position in the GTP- and GDP-bound states are shown below. **d,** Histogram of scores colored by bin (Methods). Note that 37 of the STOP mutants are toxic/GOF. **e,** Distribution of fitness scores ordered by Gsp1 sequence position, colored by mutation type: WT synonymous mutations (green), STOP codon mutations (black), and substitutions (gray).

Here we introduce an approach to generate a complete allosteric map of the essential eukaryotic GTPase switch Gsp1/Ran in the native context of its *in vivo* interaction network in *S. cerevisiae* based on comprehensive mutational perturbation^15,16^. Gsp1/Ran uses a single pair of regulators, the GAP Rna1 and the GEF Srm1, but an extended network of adaptor and effector proteins, whose interactions with Gsp1/Ran are dependent on switch state, control diverse processes including nucleocytoplasmic transport, cell cycle progression and RNA processing^6^. Gsp1 is highly conserved, with 82% of its amino acid sequence identical to the human homolog Ran. With some notable exceptions^5,17^, prior mutational scanning experiments have revealed a tolerance to mutations even among highly conserved proteins^18^, suggesting missing biological context^17,19^. In contrast, for Gsp1 in its physiological network, here we report that cellular function is affected by mutations at a large number of previously uncharacterized positions outside the active site, identifying widespread sensitivity of a central GTPase to allosteric regulation.

### Comprehensive mutational perturbation of Gsp1

To systematically measure the effect of all Gsp1 mutations on cellular function (**Fig. 1a**), we developed an approach derived from our EMPIRIC (extremely methodical and parallel investigation of randomized individual codons) method^20^ but with a generalizable plasmid dropout selection to probe the function of essential genes (**Fig. 1b**, Methods). We transformed a chromosomal GSP1 knockout strain with the wild-type (WT) GSP1 allele under the control of its native promoter on a URA selectable plasmid harboring constitutively expressed GFP, and confirmed Gsp1 protein expression via Western blot (**Supplementary File 1 Fig. 1**). We introduced a library of all possible single Gsp1 mutants, also expressed from the native Gsp1 promoter, using a HIS selectable plasmid harboring constitutively expressed mCherry. We sorted for cells expressing mCherry (library plasmid) but not GFP (WT plasmid) and compared allele abundances from the initial population to the population after six generations of growth to compute fitness scores for all 19 possible single amino acid substitutions as well as WT synonymous (WT-syn) and STOP codons at every position in Gsp1 (**Fig. 1c**, Methods). This approach interrogates variant fitness both in the presence and absence of a WT copy with the potential to inform on both gain of toxic function and loss of normal function.

We categorized the fitness score of each mutation relative to the distributions of fitness scores for WT-syn and STOP codons (**Fig. 1d**, Methods). Compared to the WT-syn distribution, 48.5% of all mutations showed deleterious fitness effects, while very few mutations (15/4315 or 0.35%) were beneficial. We observed strongly deleterious mutations in the GTPase active site, which we define as the highly conserved G1-5 functional regions of the Ras superfamily of small GTPases (including the switch loops that change conformation in the GDP- and GTP-bound states) and any additional positions contacting the nucleotide (**Fig. 1c** and **Fig. S1**).

The distribution of STOP codon scores (**Fig. 1d, e**) fell into two groups: STOP codon mutations before Gsp1 sequence position 175 had narrowly distributed fitness scores no lower than −2.90 (scores are log2-transformed changes in variant abundance relative to wild-type). In contrast, STOP mutations after position 175 had substantially lower fitness scores (down to −10.5). Residues 1-174 comprise a standard GTPase fold, whereas residues 175-219 comprise a C-terminal extension specific to the Ran subfamily (**Fig. 1c** and **Fig. S1a**). Thus, the first set of STOP codon mutants (residues 1-174) likely represent the growth defect of a null Gsp1 mutant, as internal truncations in the GTPase fold likely result in nonfunctional proteins. Mutations with worse scores than null alleles must have a functional effect more detrimental than loss-of-function, and we termed these mutations “toxic gain-of-function”, or toxic/GOF. Using a conservative definition of scores worse than the mean STOP codon mutation score of positions 1-174 by more than three standard deviations, more than half of all deleterious mutations (58.4%, and 28.4% of all mutations) were toxic/GOF. Toxic/GOF mutations were not exclusive to the active site regions defined above, but were broadly distributed across the Gsp1 structure, including in interfaces with Gsp1 partner proteins, in parts of the Gsp1 buried core, and at surface positions outside of the interaction interfaces (**Fig. 2a**).

**Figure 2.**
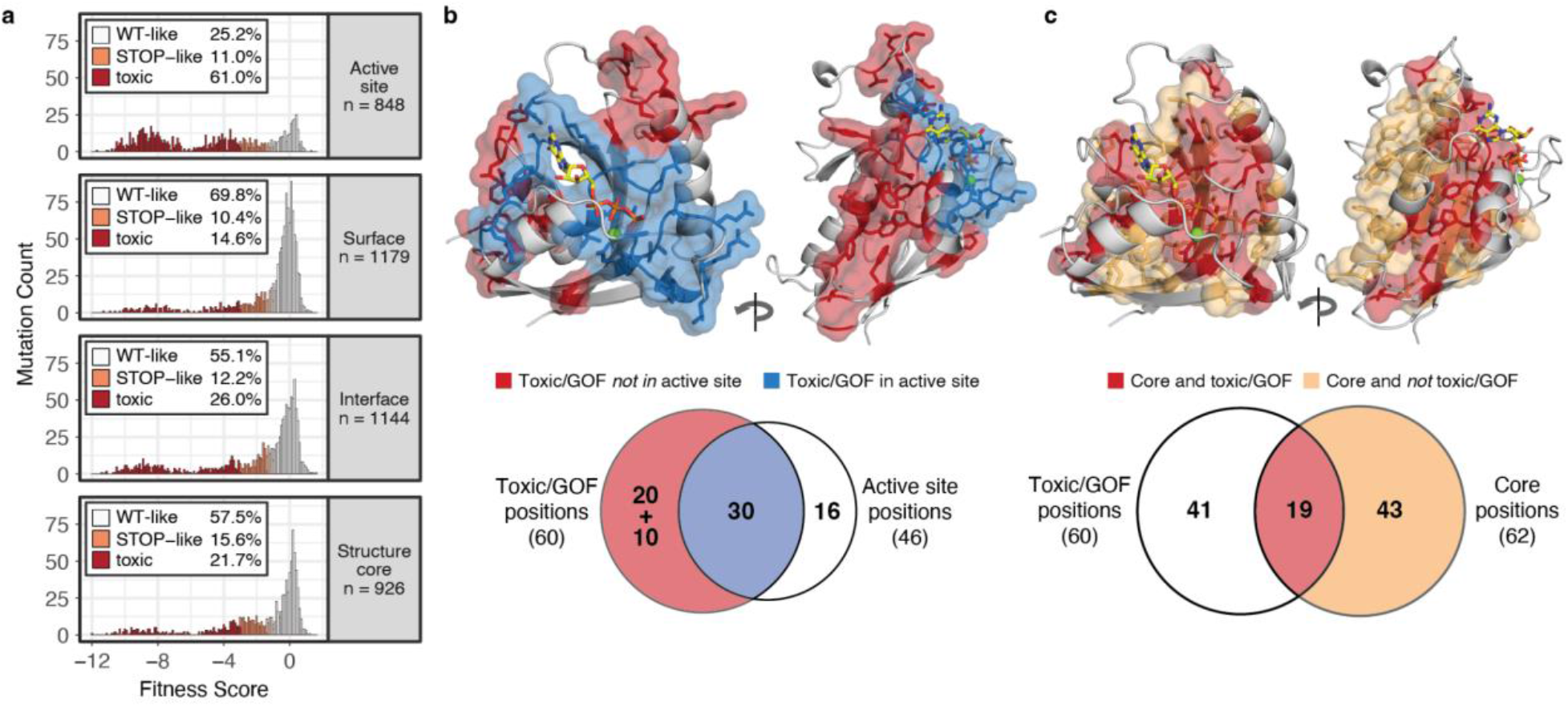
Locations of toxic/GOF positions outside of the active site. **a,** Histograms of fitness scores of mutants by structural regions; colors are as in **Fig. 1d** (showing only point mutations, excluding changes that are WT synonymous or to/from STOP; intermediate and beneficial mutations make up the difference to 100%). Fractions are computed within each structural region; n indicates number of mutations. **b** and **c,** Two views rotated by 180 degrees of the Gsp1-GTP structure (PDB ID: 3M1I) showing toxic/GOF positions in stick and surface representation (excluding the C-terminal extension). **b,** Toxic/GOF positions in the GTPase active site shown in blue, other toxic/GOF positions shown in red. Venn diagram below shows overlap of toxic/GOF positions with GTPase active site positions (10 toxic/GOF positions not shown in the structure are in the C-terminal extension). **c,** Toxic/GOF core positions shown in red, non-toxic/GOF core positions shown in orange. Venn diagram below shows overlap of toxic/GOF positions with structure core positions.

### Mapping structural locations of toxic/GOF mutations

Both the prevalence of toxic/GOF mutations and their locations across the GTPase fold were unexpected. To identify potential mechanisms underlying these findings, we defined sequence positions that were enriched in toxic/GOF mutations. We counted the number of toxic/GOF mutations at each position and compared this empirical distribution to a null distribution parameterized according to the total number of toxic/GOF mutations in the dataset (**Fig. S2**, Methods). Positions with 10 or more toxic/GOF mutations showed significant enrichment and were labeled as toxic/GOF positions. In total, 60 out of 219 Gsp1 sequence positions were toxic/GOF; 57 of these residues were identical in amino acid identity between *S. cerevisiae* Gsp1 and human Ran.

Given most substantial fitness effects observed in mutational perturbation studies are typically from mutations at positions in active sites required for function, or at positions in the protein core critical for stability, we asked whether the locations of toxic/GOF positions overlapped with the active site or the core. Only half (30/60) of the toxic/GOF positions are in the active site (**Fig. 2b**, blue) and an additional 10 positions are in the C-terminal extension. Thus, 20/60 toxic/GOF positions are at positions in the GTPase fold but distal to the active site (**Fig. 2b, red**). 16 out of the 46 active site positions are not toxic/GOF. Conversely, only 19 out of the 60 toxic/GOF positions are in the buried protein core (**Fig. 2c**, red), and 43 out of the 62 core positions are not toxic/GOF (**Fig. 2b**, orange). Moreover, mutations in the active site would typically be expected to ablate function and therefore lead to a loss-of-function phenotype (similar to STOP). However, we observe 517 (61%) toxic/GOF mutations in the active site compared to only 93 (11%) STOP-like mutations (**Fig. 2a**). Similarly, mutations in the protein core that destabilize Gsp1 would be expected to exhibit a fitness cost similar to that observed for STOP codons in the GTPase fold, but not be toxic/GOF. In addition, computational stability calculations (Methods) showed little correlation between predicted destabilization and decreased fitness when including toxic/GOF mutations, and only a modest correlation for mutations in the buried core when excluding toxic/GOF mutations (**Fig. S3, Supplementary File 1 Fig. 2**). Thus, the mechanism of Gsp1 toxic/GOF mutations is not satisfactorily explained by either simply the location in the active site or by destabilization of the protein.

### Functional roles of toxic/GOF mutants

The prevalence of toxic/GOF mutations in the C-terminal extension (**Fig. 1c, e**) provided the first evidence that the toxicity of the mutants stems from perturbed regulation: Deleting the C-terminus of Ran/Gsp1 is known to alter the balance between the switch states by stabilizing the GTP-bound form^21^, which may explain the enrichment of cancer mutations in the C-terminus of Ran^22^. We therefore asked whether all toxic/GOF mutations (**Fig. 4a**) perturbed Gsp1 GTPase switch function. This model would account for the toxic/GOF effects of mutations at the 40 positions in the GTPase active site or C-terminus. Of the remaining 20 toxic/GOF distal sites within the GTPase fold (**Fig. 2b**), 13 are located in the interfaces with key regulators of the GTPase switch Rna1 (GAP), Srm1 (GEF), and Yrb1; Y157 is an allosteric site previously identified to be coupled to the Gsp1 active site^6^, consistent with the proposed effect of mutations on regulated switching; and S155 is a known phosphorylation site^23^ neighboring the conserved G5 SAK motif in the active site (**Fig. 1c**). Four of the remaining five toxic/GOF positions are clustered in the Gsp1 structure outside of the active site, and along with the final position (H50) and two other toxic/GOF positions (N156 and F159) form distal interaction networks in crystal structures of Ran/Gsp1 that extend up to 16Å away from the nucleotide ligand to the Switch I and the C-terminal extension in the GDP-bound state^24^ (**Fig. 3a**). We verified that toxic/GOF mutants at these positions indeed had severe fitness defects compared to WT or an internal STOP-codon mutant when co-expressed with WT using a yeast spotting assay (**Fig. 3b**), and that a C-terminal deletion variant was as toxic as the toxic/GOF mutations at these positions.

**Figure 3.**
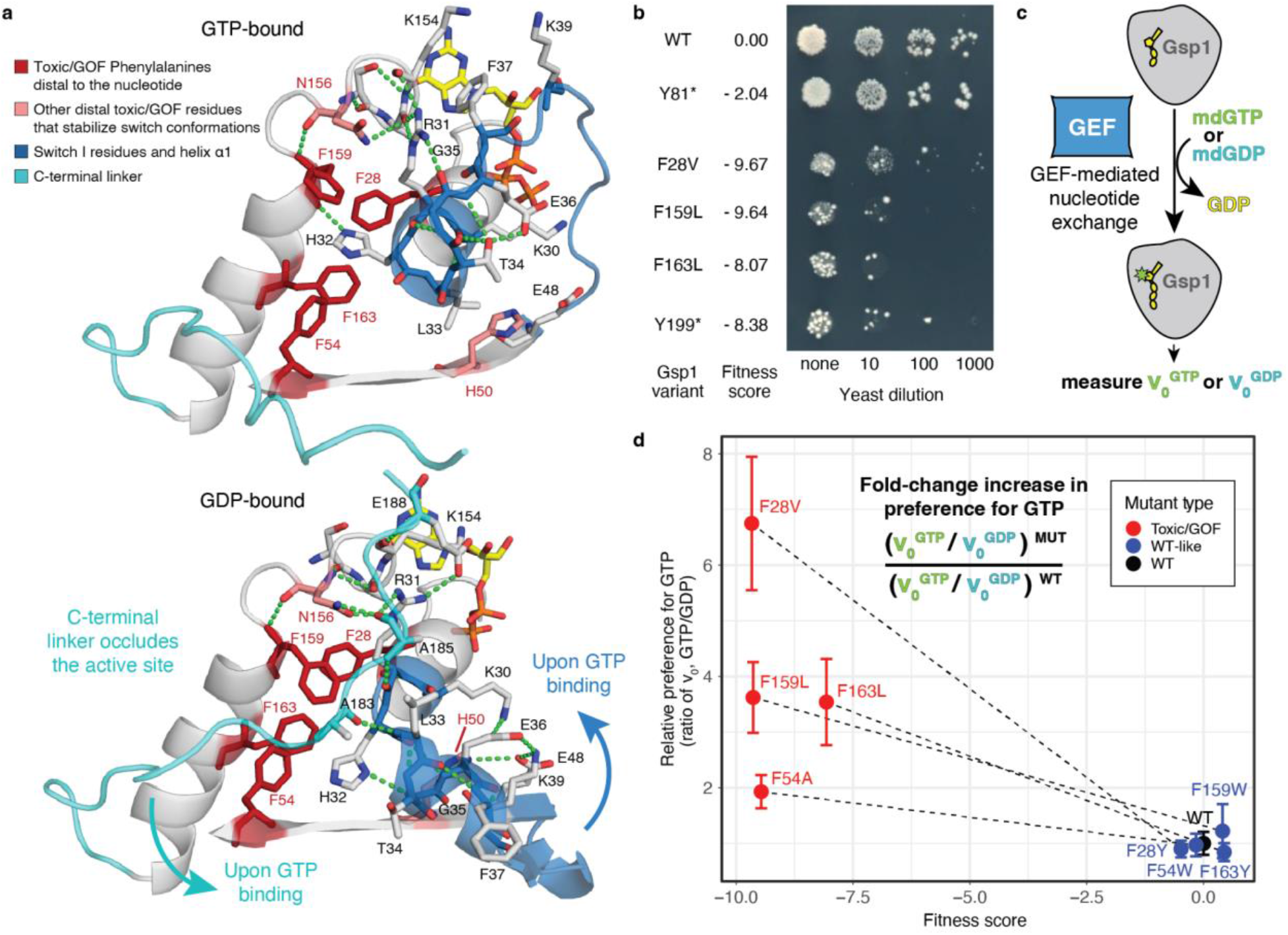
Distal toxic/GOF mutations allosterically alter the balance of the switch states. **a,** Structural depiction of extended networks of interactions in the GTP-bound (top, PDB ID: 3M1I) and GDP-bound states (bottom, PDB ID: 3GJ0). Toxic/GOF mutants characterized in panels (b) and (d) shown in red. Backbone is colored for the Switch I region (blue) and the C-terminal linker (cyan). The nucleotides are shown in yellow sticks. **b,** Plate growth assay showing a dilution series of individual Gsp1 variants expressed together with WT in *S. cerevisiae*, with corresponding fitness scores from the EMPIRIC assay. **c,** FRET-based nucleotide exchange kinetics are measured by adding an excess of mant-labeled fluorescent nucleotide and catalytic amounts of GEF to purified Gsp1 bound to GDP (Methods). **d,** Relative change in nucleotide preference for pairs of toxic and wild-type like variants at the Phe positions highlighted in (a), calculated as the ratio of initial rate of exchange to GTP divided by the initial rate of exchange to GDP, normalized to the wild-type ratio. Error bars represent the standard deviations of v0 measurements propagated across the division operator (Methods).

**Figure 4.**
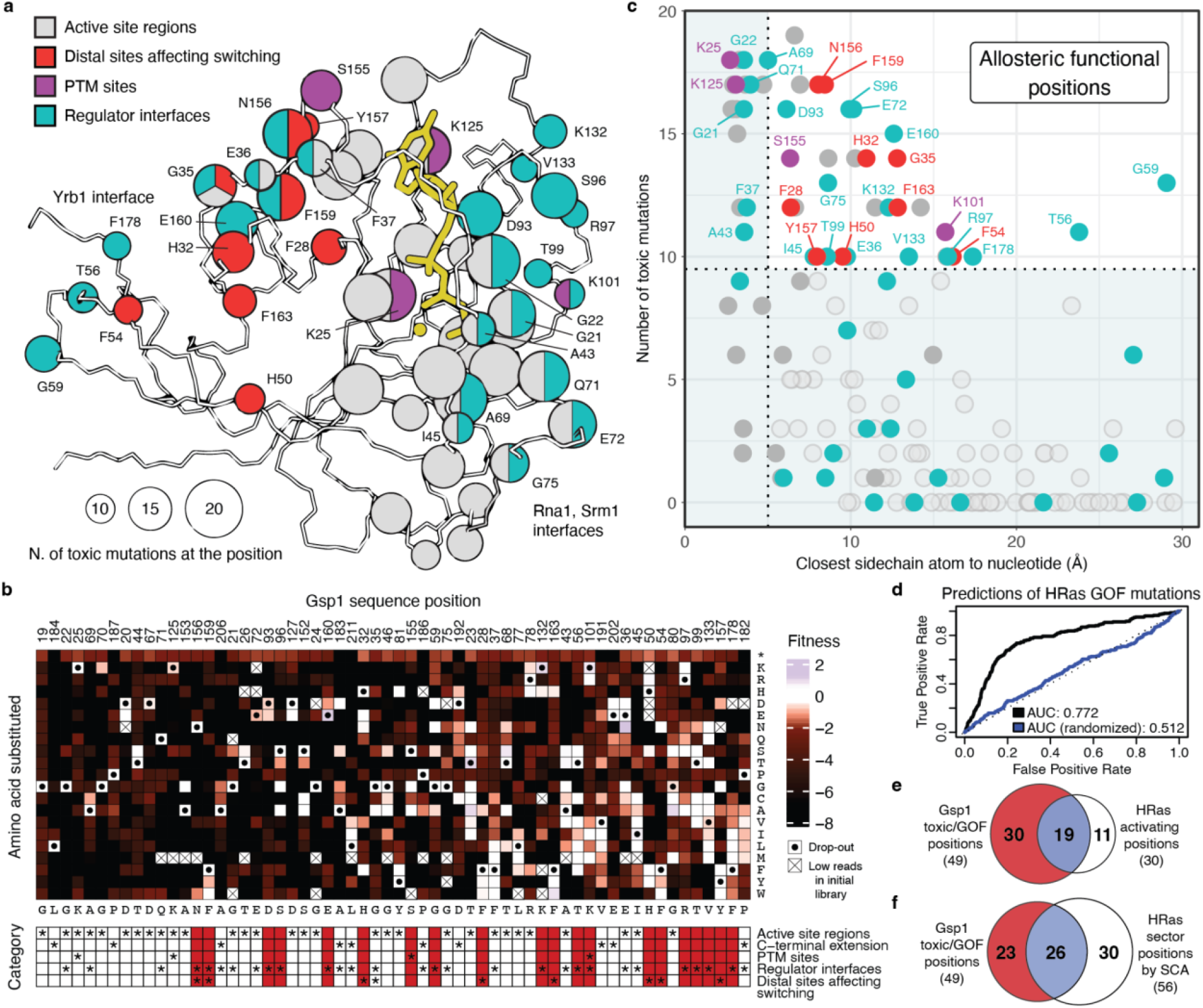
Allosteric map of the Gsp1 GTPase switch. **a,** Wire representation of Gsp1-GTP (PDB ID: 3M1I, residues 1-180). Toxic/GOF positions are shown in sphere representation. Sphere radius represents number of toxic/GOF mutations at each position. Spheres are colored by functional categories, see (b). The nucleotide and Mg^2+^cofactor are shown in yellow. **b,** Heatmap showing fitness scores at toxic/GOF positions ordered by number of toxic/GOF mutations. WT amino acid residue shown below each column. Functional annotations (stars) are shown below and marked in red for positions outside of the active site. **c,** Distance of closest sidechain heavy atom at each position to the nucleotide (GTP). Colors are as in (a). Residues not belonging to one of the four categories indicated by an open circle. **d,** Receiver operating characteristic (ROC) curves and area under the curve (AUC) showing the statistical power of Gsp1 fitness scores in classifying an *H. sapiens* HRas mutant as activating, as defined by^4^. Datasets were trimmed to the 156 sequence positions alignable for Gsp1 and HRas (**Fig. S7**). **e** and **f,** Overlap of functional sites defined as Gsp1 toxic/GOF and either (e) HRas activating or (f) comprising an HRas sector defined by statistical coupling analysis (SCA)^27^ (**Supplementary File 1 Table 1**).

To examine whether toxic/GOF mutations perturbed switch function in this unexplained set of mutants, we purified and characterized pairs of toxic/GOF (F28V, F54A, F159L, and F163L) and WT-like mutants (F28Y, F54W, F159W, and F163Y) at the four Phenylalanine positions that are clustered in the structure but distal from the active site. All purified mutants were well-folded and stable (**Extended Data Figs. 4** and **5)**. We then assessed switching by measuring the rate of GEF-mediated nucleotide exchange to either GTP or GDP using recombinantly expressed and purified *S. cerevisiae* Srm1, the GEF of Gsp1 (**Fig. 3c, Fig. S6a** and Methods). All mutants except F159L had reduced or similar GEF-catalyzed nucleotide exchange rates compared to WT (**Fig. S6b**). However, the exchange was dependent on the nucleotide: toxic/GOF mutants had a faster rate of exchange to GTP than to GDP while the WT-like counterparts had a preference for GDP over GTP, identical to WT (**Fig. 3d, Fig. S6a-c**). Hence, toxic/GOF mutations reversed the nucleotide preference of the switch but WT-like mutations did not. We also measured GAP-catalyzed GTP hydrolysis and found that toxic/GOF mutations did not have reduced GTP hydrolysis (**Fig. S6d**). We conclude that toxic/GOF mutations distal to the active site can indeed allosterically perturb the molecular function of the switch by disfavoring the GDP-bound state, while WT-like mutations at the same positions do not.

### An allosteric map of a GTPase switch

Our analyses assign functional roles to all 60 toxic/GOF positions in our dataset, mapping the functionally essential residues in a GTPase molecular switch (**Fig. 4a, b**). While the active site (nucleotide recognition sites and the GTPase switch loops) is the most common location for toxic/GOF positions, 33% of toxic/GOF positions (20/60) are outside of the active site (**Fig. 4a**). These sites are at least 5 and up to 30Å away from the nucleotide (**Fig. 4c**), showing that our method quantifying perturbations to cellular function in the native network identifies many non-local sites of allosteric regulation, even surpassing a recent study of allostery quantifying effects on biophysical function in peptide binding domains^2^.

We identify several mechanisms for how perturbations at regions outside of the active site allosterically affect GTPase switching: First, 13 sites are in interaction interfaces with the key regulators Rna1 (GAP) and Srm1 (GEF), which accelerate interconversion between the GTP-and GDP-bound states, and Yrb1, the *S. cerevisiae* homolog of human RanBP1, which stabilizes the GTP-bound state of Gsp1 and increases interaction with the GAP^21^. Second, distal positions in protein-protein interaction interfaces are in addition directly coupled to the switch by modulating the efficiency of GTP hydrolysis^6^. Third, we show here that a previously unknown allosteric cluster in the structure core (**Fig. 4a and c**, red) is coupled to switch regulation by altering the nucleotide preference (**Fig. 3**). Finally, the toxic/GOF positions also include 4 locations of posttranslational modifications (PTMs)^23,25,26^. Relatively small perturbations at all identified sites resulted in cellular defects consistently worse than a null mutant, which suggests that the effect on the rates of regulated switching between GTPase states is the key quantitative parameter dominating the functional effects of any Gsp1 mutation.

While there are no experimental studies probing the function of other GTPases under normal cellular conditions at the residue level, our functional map of Gsp1 is predictive of many activating mutations recently reported for the human H-Ras protein in mouse-derived Ba/F3 cells^4^ (**Fig. 4d**). 19/30 positions with activating mutations in H-Ras are also toxic/GOF positions in Gsp1 (**Fig. 4e**). Those positions are enriched in the active site (**Fig. S8**), whereas our Gsp1 perturbation analysis revealed additional allosteric sites including many in regulatory partner interfaces. The additional sites may be specific to Gsp1 or may not be detectable using the overactivation phenotype screened for in the H-Ras assay. Conversely, of the 11/30 activating positions not classified by our stringent cutoff as toxic/GOF in Gsp1, six have at least five toxic/GOF mutations in Gsp1, and all have at least one (**Fig. S8**). We also compared our data to a computational analysis of GTPases based on residue-residue co-variation in multiple sequence alignments of the GTPase superfamily^27^ (**Fig. 4f**). Key “sector” positions identified computationally show more overlap with the Gsp1 toxic/GOF positions than the H-Ras activation data (26/49 of the alignable positions, versus 19/49), again primarily by capturing more residues in the GTPase active site regions (**Fig. S8)**. Of the additional 30 positions suggested by the sector analysis, 12 have at least five toxic/GOF mutations in the Gsp1 data, and only four have no toxic/GOF mutations (**Fig. S8**). However, the computational sector analysis misses 23/49 toxic/GOF positions in Gsp1. This finding could indicate a lack of sensitivity or the potential for key regulatory differences between highly conserved GTPases that may be difficult to discern from sequence analysis alone but which are enabled by quantitative perturbations in the native cellular context using our approach.

### Discussion

A key finding of our work is the broad sensitivity of a critical molecular switch to perturbations at many allosteric regulatory sites outside the typically studied active site “switch” regions (**Fig. 4a, c**). We propose a model where this sensitivity of the switch facilitates both its responsiveness to many biological inputs and its output signaling specificity^6^. We identify an altered switch balance as the common mechanism by which toxic/GOF mutations affect the cellular function of Gsp1. This finding suggests that the GTPase switch balance is finely tuned and that the sensitivity of this balance to mutations at many positions might explain why GTPases are so highly conserved even outside the active site regions. We further show that relatively small perturbations to the switch balance have deleterious functional consequences. This finding is consistent with results from kinetic models of ultrasensitivity, where for switches controlled by opposing regulators (**Fig. 1a**) small changes in the concentration or activity of regulators can result in sharp changes in the fraction of the switch “on” state^28^. Our study provides an important link between allosteric regulation of the switch balance^28^ at the molecular level, and the ultrasensitivity of switches^28^ and functional consequences for cellular regulation at the systems level^6^. Our residue-level functional map of a GTPase molecular switch and the discovery of new regulatory sites opens avenues to interrogate and target GTPases controlling many essential biological processes including intracellular transport, cell growth, differentiation, and cell survival.

## Methods

### Plasmid and strain construction

To facilitate rapid Fluorescence Activated Cell Sorting (FACS)-based isolation of yeast harboring mutant Gsp1 variants, we generated plasmids marked with GFP or mCherry along with auxotrophic markers. To mimic endogenous expression of Gsp1, we cloned the Gsp1 coding sequence along with its natural promoter sequence (420 bases upstream of the start codon) and its natural 3’ region (220 bases downstream from the stop codon). We used centromeric plasmids to approximate genomic copy level. To generate a strong fluorescent signal, we used the Tef1 promoter to drive either GFP or mCherry. We cloned this Gsp1 construct into a URA-marked plasmid with GFP (pRS416Gsp1GFP), and a HIS-marked plasmid with mCherry (pRS413Gsp1mCherry).

We engineered a systematic library including all possible single amino acid changes in Gsp1 as previously described^31^. Briefly, we cloned the Gsp1 open reading frame into pRNDM and created a set of constructs with tiled inverted BsaI restriction sites bracketing 10 amino acid regions of Gsp1. For each amino acid in Gsp1, we used complementary oligonucleotides with single codons randomized as NNN to generate a comprehensive library of variants encoding all possible amino acid changes. We used Gibson assembly to transfer the library into the plasmid swap vector, generating pRS413Gsp1libmCherry. To enable library transfer, this destination vector was modified to harbor a cassette containing an SphI site along with upstream and downstream homologous sequences to Gsp1 promoter and terminator regions respectively. To facilitate short-read estimates of variant frequency we implemented a barcoding strategy as previously described^31^. We used cassette ligation at NotI and AscI restriction sites downstream of Gsp1 gene to introduce an oligonucleotide cassette including an N_18_ random sequence into the pRS413Gsp1libmCherry variants. We used paired-end Illumina sequencing to associate the 18 base barcodes with the encoded Gsp1 variants.

To generate the plasmid swap strain, DBY681, we started with a heterozygous diploid Gsp1 knockout (BY4743 Gsp1::KanMX) ordered from GE. First, we introduced pRS416Gsp1GFP and selected for transformants on synthetic media lacking uracil. Next, we sporulated the diploid transformants in order to generate haploids bearing the URA-marked plasmid. Successful transformation was evident because the selected haploid yeast cells grew on synthetic media lacking uracil, expressed GFP, grew on G418 antibiotic that selects for endogenous Gsp1 knockout, and lacked growth on synthetic media having 5-FOA which negatively selects yeast cells with URA-marked plasmid. The resulting DBY681 strain was used for all Gsp1 plasmid swap experiments.

### Gsp1 fitness competition

The DBY681 strain was made competent using the lithium acetate method^32^ and transformed with the barcoded pRS413Gsp1libmCherry plasmids. Transformation efficiency was determined by plating a small fraction of cells on selection media (SD-Ura-His+G418), aiming for five-fold coverage of the library. Sufficient transformations were performed to introduce each barcoded plasmid variant into more than 10 independent yeast cells. Following transformation, the cells were allowed to recover in synthetic dextrose media lacking uracil (SD-Ura) for ~10 hours at room temperature. The cells were then collected by centrifugation at 5000 x g for 5 minutes, washed multiple times to eliminate residual extracellular plasmid and resuspended in synthetic dextrose media lacking uracil and histidine (SD-Ura-His+G418). Sufficient media was used to achieve an optical density of approximately 0.1 at 600 nm. The cells were grown on an orbital shaker at 30 °C in the double selection media for approximately 42 hours, with constant dilution to maintain the cells in log phase.

A sample of these “initial” cells were retained for sequencing and the remainder were collected by centrifugation at 5000 x g for 10 minutes and resuspended in synthetic dextrose media lacking only histidine (SD-His) to enable loss of the URA-marked WT Gsp1 plasmid. Cells were grown in this medium with orbital shaking at 30 °C for 16 hours, which represents 6 doubling times of the parental DBY681 strain under these conditions. At the end of 16 hrs, cells were collected by centrifugation, then washed and diluted in 1x TBS with 1% BSA. For flow cytometry, the non-fluorescent parental strain W303 was treated as a negative control while DBY681 and W303 transformed with pRS413NoinsertmCherry plasmid were considered as GFP and mCherry positive controls. 3 million cells were analyzed by FACS. Cells that had lost the GFP-marked plasmid encoding WT Gsp1 were isolated by FACS. A total of 500,000 GFP-/mCherry+ cells were isolated by FACS as a sorted sample. The cells were isolated by centrifugation.

Deep sequencing was used to estimate the enrichment or depletion of mutants in the 16 hour sorted sample as compared to the initial sample in double selection media. The initial and sorted yeast samples were lysed using zymolyase and PCR amplified to generate samples for 100 bp Illumina sequencing of barcodes as previously described^31^. Briefly, primers were used that added sequences for identifying each sample as well as for compatibility with Illumina sequencing. Reads with low quality (PHRED score < 20) or that did not match in expected constant regions were eliminated from further analyses. The remaining reads were then parsed into initial and sorted bins and the number of reads of each amino acid mutation in each bin was tabulated. The experimental fitness of each variant was estimated as a selection coefficient based on the counts in the initial and sorted samples using WT synonyms for normalization using the following equation:

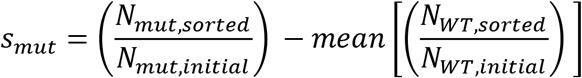

Where *s_mut_* is the selection coefficient of a mutant, *N_mut,sorted_* is the number of reads of the mutant in the sorted sample, *N_WT,sorted_* is the number of reads of WT synonyms in the sorted sample, *N_mut,initial_* is the number of reads of the mutant in the initial sample, and *N_WT,initial_* is the number of reads of a WT synonym in the initial sample. Using this equation, the average WT synonym has a selection coefficient of 0, while deleterious variants have negative s and beneficial variants have positive s. Alleles with low read counts in the initial sample, defined as less than 2% of the average variant’s number of reads, were excluded from all downstream analysis.

Fitness scores were then binned according to thresholds set by the mean and standard deviations of the distributions of scores for WT synonyms and STOP mutants. From the latter distribution we excluded mutations at sequence positions after 174, as these C-terminal STOP mutants showed significant deviations from the relatively consistent distribution of scores for STOP mutants up to and including position 174 (**Fig. 1e**) and correspond to C-terminal deletion mutants that are known to encode fully folded proteins with perturbed biochemical function^21^. Scores within two standard deviations of the mean of the WT synonym score distribution were labeled as WT-like, and scores higher than this cutoff were labeled as beneficial. For the STOP mutant distribution, scores within two standard deviations above or three standard deviations below the mean were labeled STOP-like, and scores worse than the bottom cutoff were labeled as toxic/GOF. Finally, scores between the WT-like and STOP-like distributions were labeled as intermediate.

### Expression levels of Gsp1 variants via western blot

Yeast cells were grown to exponential phase in either rich (YPD) or synthetic (SD-ura) media at 30°C. 10^8^ yeast cells were collected by centrifugation and frozen as pellets at −80°C. Cells were lysed by vortexing the thawed pellets with glass beads in lysis buffer (50 mM Tris-HCl pH 7.5, 5 mM EDTA and 10 mM PMSF), followed by addition of 2% sodium dodecyl sulfate (SDS). Lysed cells were centrifuged at 18,000 x g for 1 min to remove debris, and the protein concentration of the supernatants was determined using a BCA protein assay kit (CAT #23227, Pierce) compared to a Bovine Serum Albumin (BSA) protein standard. 25 μg of total cellular protein was resolved by SDS-PAGE and was either visualized with Coomassie blue stain, or transferred to a PVDF membrane, and probed using Rabbit anti-RAN primary (CAT # PA 1-5783, ThermoFisher Scientific) and Donkey anti-Rabbit HRP-linked secondary (CAT # NA934V, Cytiva Life Science) and visualized with ECL-2 substrate (CAT #80196, Pierce).

### Yeast spotting assays

Individual variants of Gsp1 were generated by site-directed mutagenesis using overlapping mutagenic PCR primers and confirmed by Sanger sequencing. Variants were cloned in a HIS-marked plasmid (pRS413 with mCherry). For the yeast spotting assays, the plasmids were transformed into DBY681 (Gsp1::kan, pRS416Gsp1 with GFP) using the lithium acetate method^32^. Transformed cells were recovered in SD-ura media for 6 hours and then 5 uL of a 10x dilution series of cells were spotted onto SD-ura-his plates. For the bacterial spotting assays, the same plasmids were transformed into chemically competent E. coli, recovered for 1 hour in LB, and 5 uL of a 10x dilution series of cells were spotted on LB-amp plates.

### Statistical modeling of the distribution of toxic/GOF mutations

A hypergeometric distribution was used to model the null distribution of toxic/GOF mutations partitioning among the 219 residue positions. This approach computes the probability that a certain number of toxic/GOF scores would be at the same position, given the number of toxic/GOF scores in the dataset and 21 possibilities at each position (20 amino acids and STOP). The calculation was performed using the *dhyper* function in the *stats* package of the programming language *R*.

### Protein purifications

Gsp1 variants were expressed from a pET-28 a (+) vector with an N-terminal 6xHis tag in E. coli strain BL21 (DE3) in the presence of 50 mg/L Kanamycin in autoinduction EZ medium for 60 hours at 20 °C^33^. The autoinduction medium consisted of ZY medium (10 g/L tryptone, 5 g/L yeast extract) supplemented with the following stock mixtures: 20xNPS (1M Na_2_HPO_4_, 1M KH_2_PO_4_, and 0.5 M (NH_4_)2SO_4_), 50x 5052 (25% glycerol, 2.5% glucose, and 10% α-lactose monohydrate), 1000x trace metal mixture (50 mM FeCl_3_, 20 mM CaCl_2_, 10 mM each of MnCl_2_ and ZnSO_4_, and 2 mM each of CoCl_2_, CuCl_2_, NiCl_2_, Na_2_MoO_4_, Na_2_SeO_3_, and H_3_BO_3_ in ~60 mM HCl). Cells were lysed in 50 mM Tris pH 7.5, 500 mM NaCl, 10 mM MgCl2, 10 mM imidazole, and 2 mM β-mercaptoethanol using a microfluidizer from Microfluidics. The His-tagged proteins were purified on Ni-NTA resin (Thermo Scientific #88222) and washed into a buffer of 50 mM Tris (pH 7.5), 100 mM NaCl, and 4 mM MgCl2. The N-terminal His-tag was digested at room temperature overnight using 12 NIH Units per mL of bovine thrombin (Sigma-Aldrich T4648-10KU). Proteins were then bound to an additional 1 mL of Ni-NTA resin to remove non-specific binders and passed through a 0.22 uM filter. Purity was confirmed to be at least 90% by SDS polyacrylamide gel electrophoresis. Samples were concentrated on 10 kDa spin filter columns (Amicon Catalog # UFC901024) into a storage buffer of 50 mM Tris pH 7.5, 150 mM NaCl, 4 mM MgCl2, and 1 mM Dithiothreitol. Using this protocol, Gsp1 variants are purified bound to GDP (as any bound GTP is likely hydrolyzed completely during the lengthy incubation steps beginning with thrombin cleavage). The complete hydrolysis to GDP was confirmed for this protocol previously^6^ using reverse phase high performance liquid chromatography on a C18 column. Protein concentrations were confirmed by measuring at 10-50x dilution using a Nanodrop (ThermoScientific). The extinction coefficient at 280 nm used for wild-type Gsp1 was 37675 M^-1^ cm^-1^, based on the value calculated from the primary protein sequence using the ProtParam tool (https://web.expasy.org/protparam/) accounting for the cleaved N-terminal residues, and augmented by 7765 M^-1^ cm^-1^ to account for the bound nucleotide, as described previously (see Note 4.13 by Smith and Rittinger^34^). Extinction coefficients were calculated for each Gsp1 mutant by the same method. The ratio of absorbance at 260 nm and 280 nm for purified Gsp1 bound to GDP was 0.76 for all mutants except for N156W, for which it was 1.34. Concentrated proteins were flash-frozen and stored at −80 °C.

*S. cerevisiae* Srm1 (GEF, Uniprot P21827) and *S. pombe* Rna1 (GAP, Uniprot P41391) were also expressed from a pET-28 a (+) vector with a N-terminal 6xHis tag in *E. coli* strain BL21 (DE3). For discussion on the appropriateness of using *S. pombe* GAP for kinetics studies of *S. cerevisiae* Gsp1, see the Supplementary Discussion of ^6^. Srm1 was purified as Δ1-27Srm1 and GAP as a full-length protein. ScΔ1-27Srm1 and SpRna1 were expressed in 2xYT medium (10 g NaCl, 10 g yeast extract (BD BactoTMYeast Extract #212720), 16 g tryptone (Fisher, BP1421) per 1 L of medium) in the presence of 50 mg/L Kanamycin overnight at 25 °C. Expression was induced by addition of 300 μmol/L Isopropyl-β-D-thiogalactoside (IPTG). Cells were lysed in 50 mM Tris pH 7.5, 500 mM NaCl, 10 mM imidazole, and 2 mM β-mercaptoethanol using a microfluidizer from Microfluidics. The His-tagged proteins were purified on Ni-NTA resin (Thermo Scientific #88222) and washed into a buffer of 50 mM Tris (pH 7.5) and 100 mM NaCl. The N-terminal His-tag was digested at room temperature overnight using 12 NIH Units per mL of bovine thrombin (Sigma-Aldrich T4648-10KU). Proteins were then bound to an additional 1 mL of Ni-NTA resin to remove non-specific binders and passed through a 0.22 uM filter. Proteins were then purified using size exclusion chromatography (HiLoad 26/600 Superdex 200 pg column from GE Healthcare), and purity was confirmed to be at least 90% by SDS polyacrylamide gel electrophoresis. Samples were concentrated on 10 kDa spin filter columns (Amicon Catalog # UFC901024) into storage buffer (50 mM Tris pH 7.5, 150 mM NaCl, 1 mM Dithiothreitol). Protein concentrations were confirmed by measuring at 10-50x dilution using a Nanodrop (ThermoScientific). Extinction coefficients were estimated based on their primary protein sequence using the ProtParam tool (https://web.expasy.org/protparam/). Concentrated proteins were flash-frozen and stored at −80 °C.

### Circular dichroism (CD) spectroscopy

Samples for CD analysis were prepared to a concentration of 1 - 2.5 μM Gsp1 in 2.5 mM Tris pH 7.5, 5 mM NaCl, 200 μM MgCl_2_, and 50 μM Dithiothreitol. CD spectra were recorded at 25 °C using 1- or 2-mm cuvettes (Starna, 21-Q-1 or 21-Q-2) in a JASCO J-710 CD-spectrometer (Serial #9079119). The bandwidth was 2 nm, rate of scanning 20 nm/min, data pitch 0.2 nm, and response time 8 s. Each CD spectrum represents the accumulation of 5 scans. Buffer spectra were subtracted from the sample spectra using the Spectra Manager software Version 1.53.01 from JASCO Corporation. Temperature melts were performed from 25 °C - 95 °C, monitoring at 210 nm, using a data pitch of 0.5°C and a temperature slope of 1°C per minute. As all thermal melts of wild-type and mutant Gsp1 proteins were irreversible, only apparent *T_m_* was estimated by fitting melts to a two-state unfolding equation:

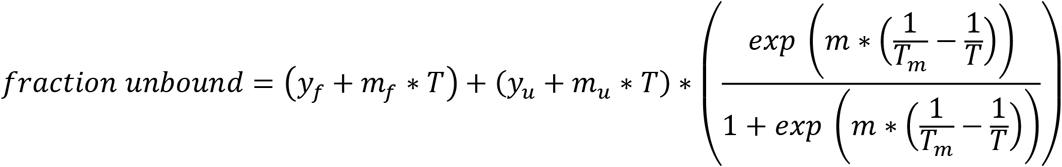

with *T* corresponding to the temperature in degrees Celsius, *y_u_* and *y_f_* corresponding to the molar ellipticity signal at the unfolded and folded states, and *m_u_*, *m_f_*, and *m* corresponding to the slopes of signal change at the unfolded state, the folded state, and the state transition.

### Kinetic measurements of GEF-mediated nucleotide exchange

Kinetic parameters of GEF mediated nucleotide exchange were determined using a fluorescence resonance energy transfer (FRET) based protocol as previously described^6^. Gsp1 variants are purified as a Gsp1:GDP complex, as verified previously^6^. Nucleotide exchange from GDP to either mant-dGDP (3’ - O - (N - Methyl - anthraniloyl) - 2’ - deoxyguanosine - 5’ - diphosphate, CAT # NU-205L, Jena Biosciences) or mant-dGTP (3’ - O - (N - Methyl - anthraniloyl) - 2’ - deoxyguanosine 5’ triphosphate, CAT # NU-212L, Jena Biosciences) was monitored by measuring a decrease in intrinsic Gsp1 tryptophan fluorescence (295 nm excitation, 335 nm detection) due to FRET upon binding of the mant group. Experiments were performed in 100 μl reaction volumes containing GTPase assay buffer (40 mM HEPES pH 7.5, 100 mM NaCl, 4 mM MgCl2, 1 mM Dithiothreitol) using 5 μM Gsp1, 2.5 nM Srm1 (GEF), and 100 μM mant-labeled nucleotide. Time courses were collected for 20 min at 30°C in a Synergy H1 plate reader from BioTek, using Corning 3686 96-well half-area non-binding surface plates. Initial rates *v_0_* of nucleotide exchange were estimated using linear fits to the very beginning of reactions for all variants except F28V. Due to the especially slow exchange rate of F28V, the reactions maintained linearity over the entire time course, and so the true exchange rate was estimated by subtracting the rate of background fluorescence decay (obtained from a control without GEF in a separate well on the same plate) from a linear fit of the full time course. At least four replicates were performed for each variant, allowing for calculation of the standard deviation of *v_0_* values (*sd*). The preference for GTP over GDP was calculated as 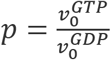 with the error of preference (e) being computed using error propagation over the division operator:

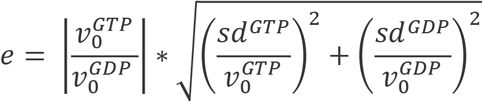

Finally, the relative change in preference 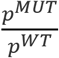 was calculated for each mutant, with the error once again propagated across the division operator. All relative changes in preference were computed using WT rates fit on the same day using the same aliquot of GEF, to normalize for any errors in enzyme concentration measurements. Furthermore, experiments for pairs of toxic/GOF and WT-like mutants were always performed on the same day using the same aliquots of GEF.

### GTP loading of Gsp1 for GAP-activated hydrolysis assay

WT Gsp1 was loaded with GTP by incubation in the presence of 20-fold excess GTP (Guanosine 5’-Triphosphate, Disodium Salt, CAT # 371701, Calbiochem) in 50 mM Tris HCl pH 7.5, 100 mM NaCl, 4 mM MgCl_2_. Exchange of GDP for GTP was initiated by the addition of 10 mM EDTA. Reactions were incubated for 3 hours at 4°C and stopped by addition of 1 M MgCl_2_ to a final concentration of 20 mM MgCl_2_ to quench the EDTA. GTP-loaded protein was buffer exchanged into a GTPase assay buffer of 40 mM HEPES pH 7.5, 100 mM NaCl, 4 mM MgCl_2_, 1 mM DTT using NAP-5 Sephadex G-25 DNA Grade columns (GE Healthcare # 17085301).

### Kinetic measurements of GAP-activated GTP hydrolysis

Kinetic parameters of the GTP hydrolysis reaction were determined as previously described^6^. Gsp1 samples for GTP hydrolysis kinetic assays were first loaded with GTP as described above. GTP hydrolysis was monitored by measuring fluorescence of the E. coli phosphate-binding protein labeled with 7 - Diethylamino - 3 - [N - (2 - maleimidoethyl) carbamoyl] coumarin (MDCC) (phosphate sensor, CAT # PV4406, Thermo Fisher) upon binding of the free phosphate GTP hydrolysis product (excitation at 425 nm, emission at 457 nm). Experiments were performed in 100 μl GTPase assay buffer (40 mM HEPES pH 7.5, 100 mM NaCl, 4 mM MgCl_2_, 1 mM Dithiothreitol) using 5 μM Gsp1:GTP, 1 nM *Sp*Rna1 (GAP), and 20 μM phosphate sensor. Time courses were collected for 60 min at 30°C in a Synergy H1 plate reader from BioTek, using Corning 3881 96-well half-area clear-bottom non-binding surface plates. A conversion factor between fluorescence and phosphate concentration was calibrated for the 20 μM concentration of the sensor with a range of concentrations of K_2_HPO_4_, considering only data in the linear range. For each individual GAP-mediated GTP hydrolysis experiment, a control experiment with the same concentration of GTP-loaded Gsp1 and the same concentration of sensor, but without added GAP, was run in parallel. The first 100 s of these data were used to determine the baseline fluorescence. The kinetic parameters (*k_cat_* and *K_m_*) were estimated by directly analyzing the full reaction progress curve with an analytical solution of the integrated Michaelis-Menten equation, as done previously^6^ using the custom-made software DELA^35^. Specifically, each time course was fitted to an integrated Michaelis Menten equation:

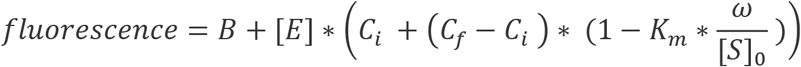

Where [*E*] is the total enzyme (GAP) concentration, *C_i_* is the initial fluorescence, *C_f_* is the final fluorescence, [*S*]_0_ is the initial concentration of the substrate (Gsp1:GTP), and *B* is the baseline slope in fluorescence per second. Exact concentration of loaded Gsp1:GTP [*S*]_0_ was estimated based on the plateau fluorescence and the sensor calibration parameters to convert the fluorescence to free phosphate concentration. The *ω* parameter was solved by using the Lambert *ω* algorithm,

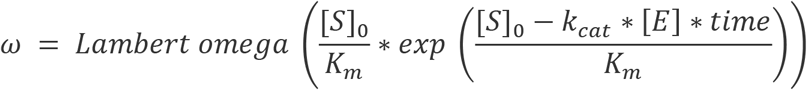

### Structural bioinformatics

Protein structures were downloaded from the PDB-REDO databank web server^36^. Secondary structure annotation of the GTP-bound (PDB 3M1I, chain A) and GDP-bound (PDB 3GJ0) states were performed using PyMOL (Schrödinger, Inc.) with the command *ss H/S*, followed by manual inspection and comparison to the results of the DSSP algorithm^37^ implemented in the PyRosetta interface (version *2020.28+release.8ecab77aa50*) to the Rosetta molecular modeling suite^38^.

Assignments of structural regions (structure core, interface core, and surface) of Gsp1 were previously reported^6^ whereby burial of a residue (in either the structure core or interface core) was defined based on per-residue relative solvent accessible surface area (rASA)^39^ compared to the empirical maximum solvent accessible surface area for each of the 20 amino acids^40^. Annotations of the canonical Ras superfamily GTPase regions were taken from^29^ as well as studies of Ran structures^30,41–43^. The key GEF binding region annotations were taken from^44^.

### Rosetta ΔΔG calculations

Stability calculations for all 19 possible point mutations were performed using the application *cartesian-ddg^45,46^* in the Rosetta software suite. Calculations were performed for both the GTP-bound (PDB 3M1I, chain A) and GDP-bound (PDB 3GJ0) structures. First, the structures were minimized in cartesian coordinates using the *relax* application, the *ref2015_cart* score function, and constraints to starting coordinates. The *relax* protocol was run 20 times and the lowest scoring structure was chosen. The GTP-bound structure was truncated after position 183, as the C-terminal extension contains unresolved regions in this crystal structure and adopts a different conformation when bound to Yrb1. The prepared starting structures were then run through the *cartesian-ddg* protocol, which computes energy scores in Rosetta Energy Units (REU) for each mutation by choosing the best scoring rotamer for the mutant amino acid, then minimizing the structure 5 times in cartesian coordinates while only allowing movement of sidechain atoms within a 6Å window around the mutated residue and backbone atoms within a three residue window (1 neighboring residue on each side), and finally taking the average score of the 5 structures. ΔΔG scores are computed by performing the same protocol at each site while choosing the best WT rotamer at the first step, and then taking the difference between the mutant and WT energies. Finally, the ΔΔG values were scaled down using a scaling factor of 0.298, determined from a benchmark set of stability calculations (performed in parallel with the Gsp1 calculations) for which experimental ΔΔG values are available^47,48^, as recommended by the authors of the *cartesian-ddg* protocol^45^. Position Q71 was excluded from the analysis, as the GTP-bound structure harbors a Q71L mutation at the catalytic glutamine. The full set of command line flags for the *relax* and *cartesian-ddg* protocols are shown below. The *movemap* file *gsp1.movemap* was not included for *relax* runs on the benchmark set. All associated configuration files as well as the datasets of Gsp1 and benchmark set ΔΔG values are available in full at the code repository at https://github.com/cjmathy/Gsp1_DMS_Manuscript.

*relax* flags:

~~~
<path/to/Rosetta>/main/source/bin/relax.default.linuxgccrelease \
-s <path/to/pdb_file> \
-out:path:all <path/to/output_dir> \
-database <path/to/Rosetta>/main/database \
-use_input_sc \
-in:file:movemap gsp1.movemap \ # for Gsp1 structures only
-constrain_relax_to_start_coords \
-ignore_unrecognized_res \
-nstruct 20 \
-relax:cartesian \
-relax:coord_constrain_sidechains \
-relax:min_type_lbfgs_armijo_nonmonotone \
-score:weights ref2015_cart \
-relax:script cart2.script
~~~

*gsp1.movemap:*

~~~
RESIDUE * BBCHI
RESIDUE 201 202 NO # Nucleotide and Mg in 3M1I. Use 208 209 for 3GJ0
JUMP * YES
~~~

*cart2.script:*

~~~
switch:cartesian
repeat 2
ramp_repack_min 0.02  0.01    1.0 50
ramp_repack_min 0.250 0.01    0.5 50
ramp_repack_min 0.550 0.01    0.0 100
ramp_repack_min 1     0.00001 0.0 200
accept_to_best
endrepeat
~~~

*cartesian-ddg* flags:

~~~
<path/to/Rosetta>/main/source/bin/cartesian_ddg.linuxgccrelease \
           -database <path/to/Rosetta>/main/database \
           -s <path/to/relaxed_pdb_file> \
           -out:path:all <path/to/output_dir> \
           -ddg:mut_file <path/to/mut_file> \
           -ddg:output_dir <path/to/output_dir> \
           -ddg:iterations 5 \
           -ddg::cartesian \
           -ddg::dump_pdbs true \
           -ddg::bbnbrs 1 \
           -fa_max_dis 9.0 \
           -score:weights ref2015_cart
~~~

Example *mut_file*, which specifies the mutation to make (here, F159L, which is residue 150 in the numbering scheme used by Rosetta, which always starts at 1 for the N-terminal residue in a chain). One such *mut_file* is used for each modeled mutation.

~~~
total 1 # specifies only one mutation is being made
1       # specifies only one mutation is being made
F 150 L
~~~

### Comparison to H-Ras mutagenesis data

Alignment of sequence positions between Gsp1 and H-Ras was performed with the *bio3d* package^49^ using the function *pdbaln* followed by refinement of the alignment upon inspection of the structural superposition using the function *pdbfit*. PDB structures used for the superposition were 3M1I, 1K5D, 1WQ1, and 3L8Z. The sequence alignment is shown in **Fig. S7**. In total, 156 structurally aligned positions were included in the analysis. Fitness scores from the human H-Ras mutagenesis study^4^ were obtained from datasets deposited on GitHub at https://github.com/fhidalgor/ras_cancer_hidalgoetal (commit *0dcb01b* from Dec. 22, 2021, downloaded on January 31, 2022). Receiver operating characteristic (ROC) curves were produced as described in the original study^4^, namely by considering H-Ras mutations with a fitness score greater than 1.5 times the standard deviation in the Ba/F3 dataset as activating (true positives), with the other mutations labeled as true negatives. Then, a variable threshold value of Gsp1 fitness is used, and for each threshold value, mutations with a Gsp1 fitness score less than that threshold (starting with the most deleterious mutations and proceeding to decreasingly deleterious Gsp1 mutations) are considered to predict H-Ras activation.

For the analysis of overlap with Gsp1 toxic/GOF positions (**Fig. 4e**, **Fig. S8**), a threshold of 2 or more activating mutations at a position was chosen for defining H-Ras activation positions, since a large number of positions have only a single activating mutation out of the 21 possible mutations. This threshold was supported by a chi-squared test evaluating the strength of association between the Gsp1 toxic/GOF and H-Ras activating sets when applying the threshold (P = 7.711 e-5) vs. including all positions with one or more activating mutation (P = 0.0411).

### Comparison to Statistical Coupling Analysis

H-Ras sector positions identified by statistical coupling analysis^50^ were taken from an analysis notebook document by the Ranganathan group publicly available on their Github (https://github.com/ranganathanlab/pySCA/blob/master/notebooks/SCA_G.ipynb, commit *301f874*, downloaded on February 9, 2022) prepared in concert with their study^27^.

## Acknowledgments

We would like to thank Mark Kelly for advice and training on biophysical experiments; David Lambright for generously providing kinetic analysis software; Frank DiMaio, Hahnbeom Park, and Brandon Frenz for advice on Rosetta ΔΔG calculations; members of the Kortemme lab for discussion; and Guy Riddihough of Life Science Editors for helpful comments and edits on the manuscript. This work was supported by a grant from the National Institutes of Health (R01-GM117189) to T.K. and D.N.A.B.; C.J.P.M. is a UCSF Discovery Fellow; T.K. is a Chan Zuckerberg Biohub Investigator.

## Author contributions

C.J.P.M., D.N.A.B. and T.K. identified and developed the core questions. C.J.P.M. performed the majority of the data analysis, with contributions from D.M., T.P., P.M., D.N.A.B. and T.K., and the biophysical experiments. P.M. performed the mutational scanning experiments. J.M.F. performed the spotting assays and western blot experiments. C.J.P.M. and T.K. wrote the manuscript with contributions from the other authors. D.N.A.B. and T.K. oversaw the project.

## Competing interest

The authors declare no competing interests.

**Supplementary Information** is available for this paper.

## Data and Code availability

All relevant data and code are available in the manuscript, the supplementary materials, or on GitHub at https://github.com/cjmathy/Gsp1_DMS_Manuscript.

---

**Figure S1.**
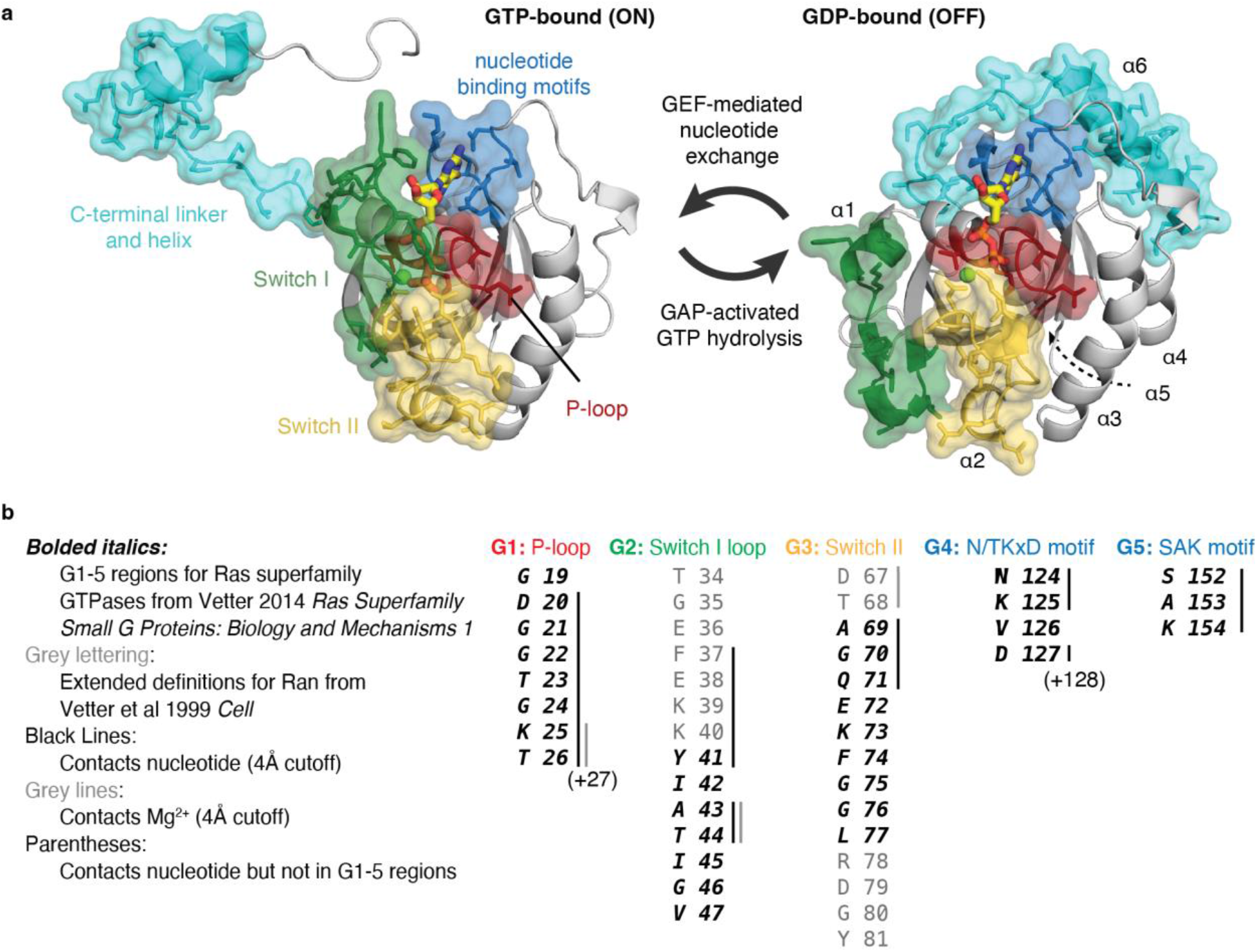
Structural annotation of Ran/Gsp1 GTPase regions. **a,** Structures of GTP-bound *S. cerevisiae* Gsp1 (PDB ID: 3M1I co-complex partners Yrb1 and Xpo1 not shown) and GDP-bound *H. sapiens* Ran (PDB ID: 3GJO). G1-5 regions corresponding to **Fig. 1b** are shown in surface representation: P-loop (red), Switch I loop (green), Switch II loop (yellow), nucleotide binding motifs (blue), C-terminal extension (cyan). Nucleotides are shown in yellow stick representation, and the Mg^2+^ cofactors are shown as green spheres. Large conformational changes associated with state-switching occur in the Switch I and II loops as well as the C-terminal extension. **b,** Residues comprising the G regions, highlighting the distinction between canonical definitions derived from evolutionary conservation analysis of all Ras superfamily GTPases^29^ and the Ran/Gsp1 specific definitions derived from structural characterization^30^. All sequence numbers shown correspond to *S. cerevisiae* Gsp1.

**Figure S2.**
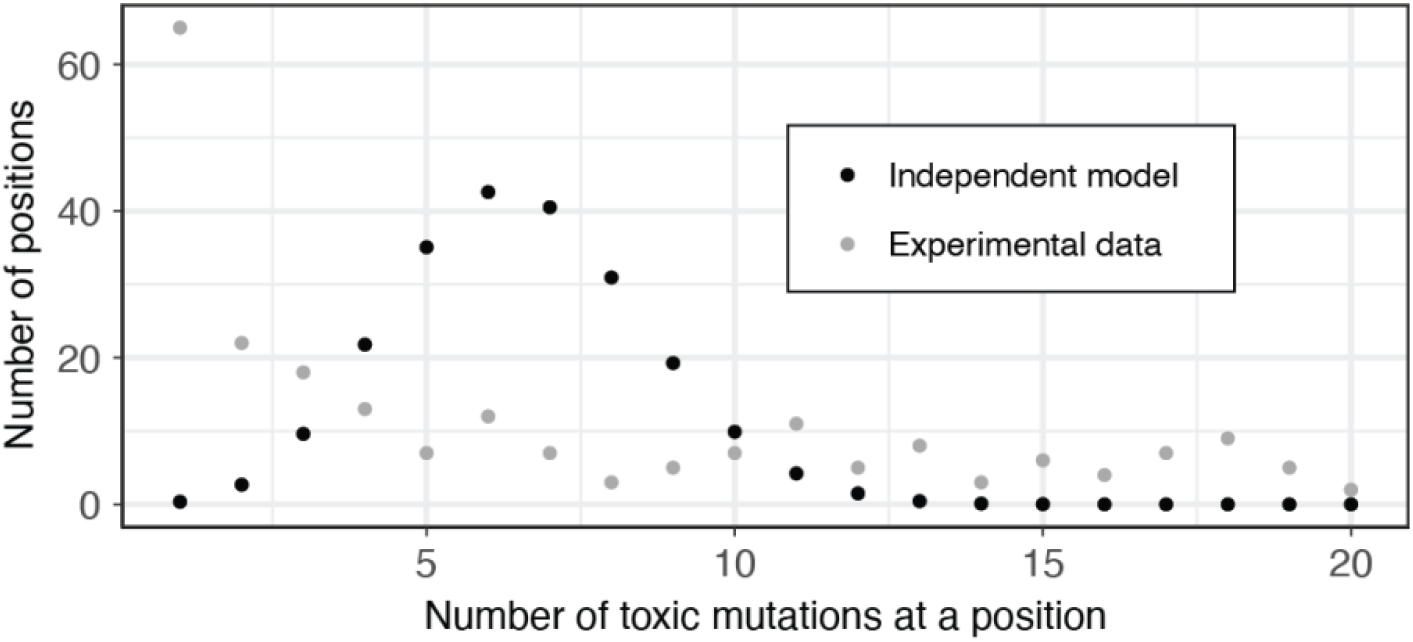
Null model of the distribution of toxic/GOF mutations used to define toxic/GOF positions. A null distribution for the number of toxic/GOF mutations observed at each of the possible 219 positions in Gsp1 was constructed from the hypergeometric distribution (Methods) and compared to the observed number of toxic/GOF mutations at each position. We chose a threshold of 10 or more toxic/GOF mutations at a position to define a position as toxic/GOF.

**Figure S3.**
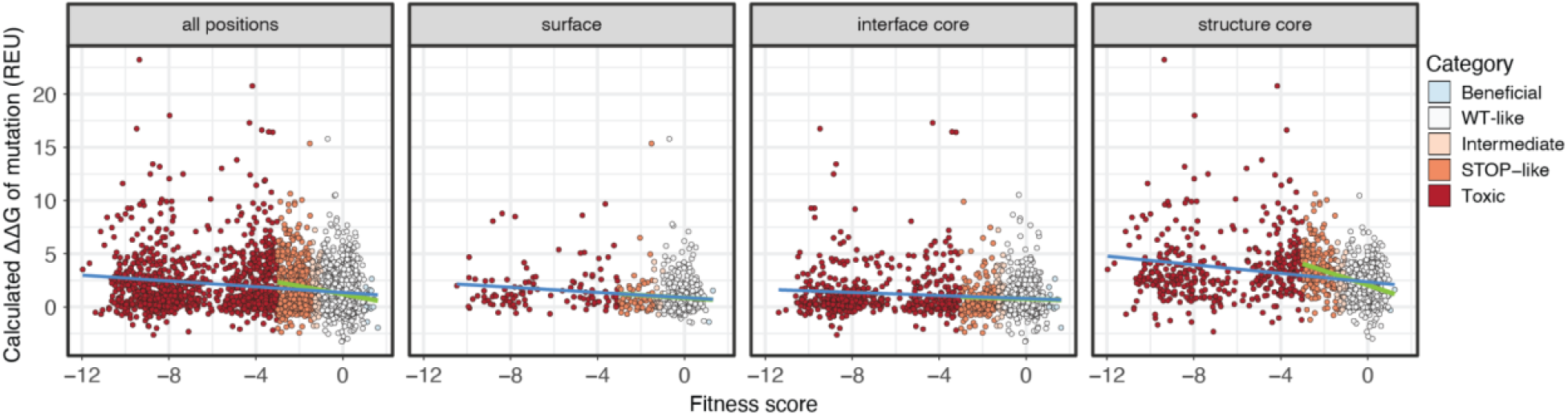
Prediction of effects of mutation on protein stability (ΔΔG) using Rosetta. Scatterplot comparing the EMPIRIC fitness score with the calculated change in protein stability upon mutation (ΔΔG in Rosetta Energy Units, REU) predicted using the GTP-bound structure (PDB ID: 3M1I, residues 10-183). Points are colored by score category as in **Fig. 1d**. Scatterplots broken down by structural region are also shown. Lines indicate best fit when including (blue) or excluding (green) the toxic/GOF mutations.

**Figure S4.**
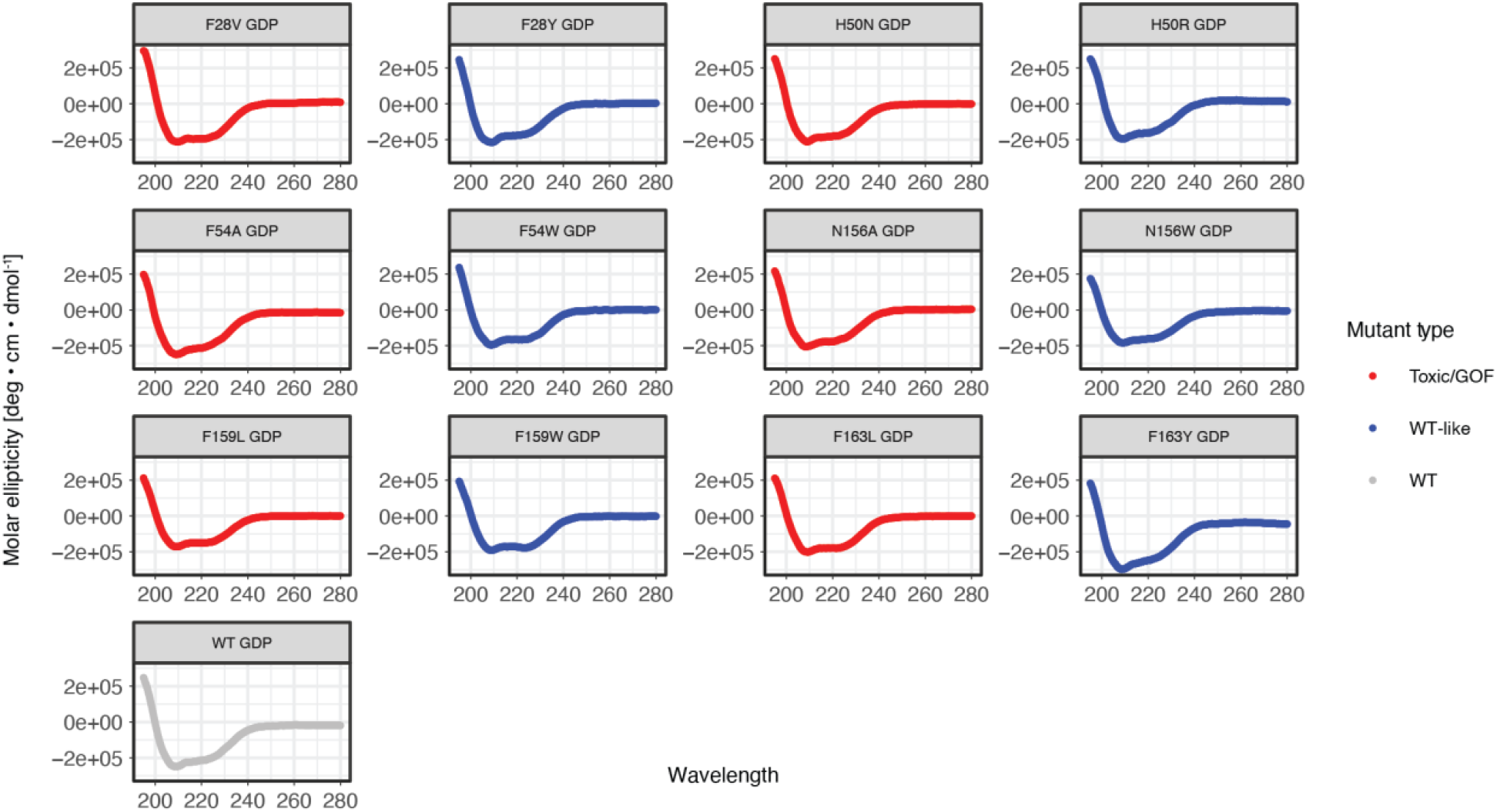
Circular dichroism (CD) spectra of purified Gsp1 variants. CD spectra for Gsp1 variants at 25°C. Variants with a toxic/GOF mutation are shown in red, variants with a WT-like mutation in blue, and WT in gray.

**Figure S5.**
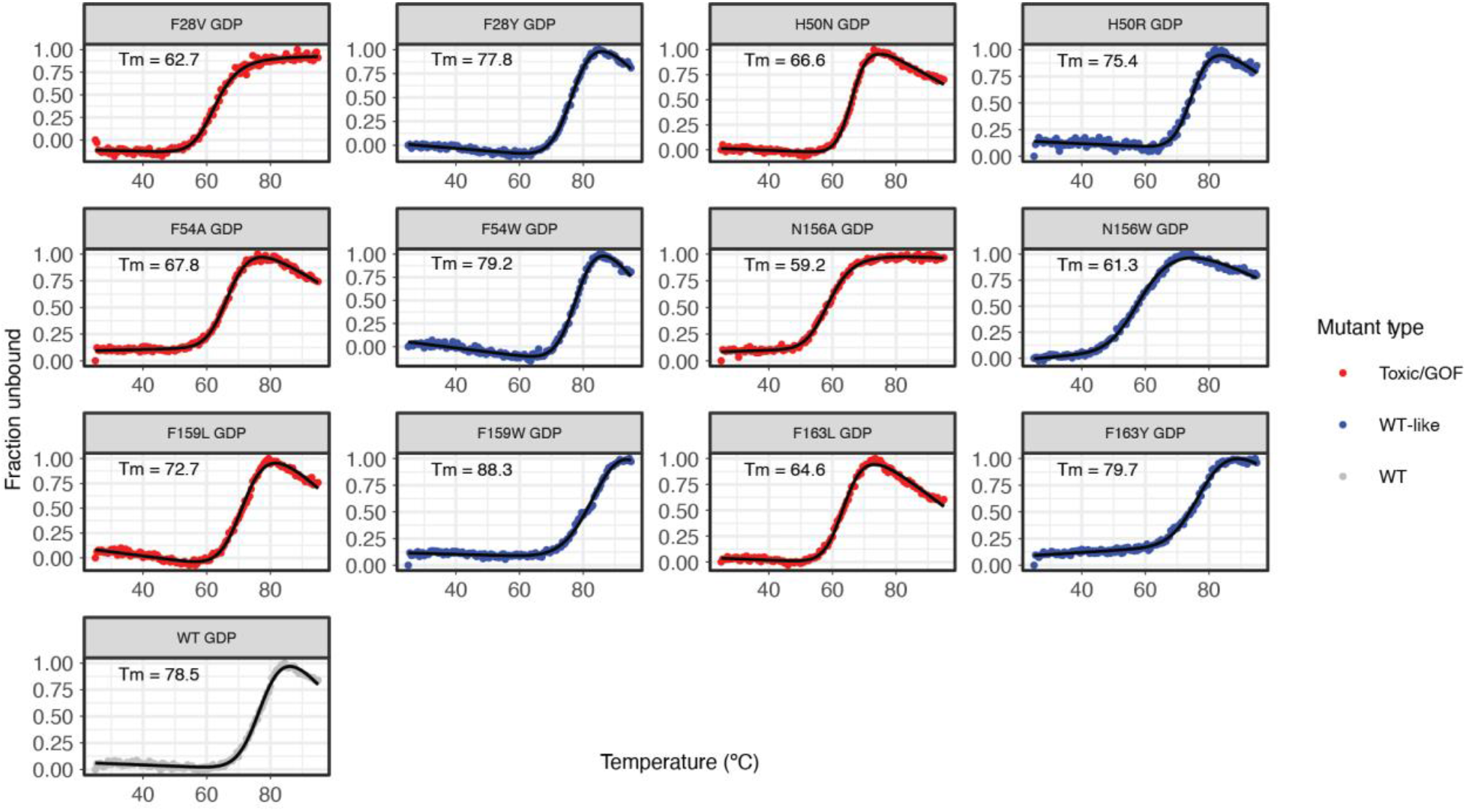
Circular dichroism (CD) thermal melts of purified Gsp1 variants. CD melts of Gsp1 variants from 25 - 95°C. Variants with a toxic/GOF mutation are shown in red, variants with a WT-like mutation in blue, and WT in gray. Apparent melting temperature 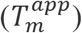 values were computed by fitting melts to a two-state unfolding equation (see Methods). CD melts are not reversible. All variants are stable up to at least 50°C, although toxic/GOF mutations resulted in slightly decreased apparent melting temperatures (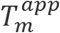 62.7–72.7°C) compared to WT (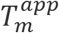 78.5°C) or WT-like mutations (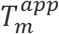 77.8–88.3°C) at the same positions.

**Figure S6.**
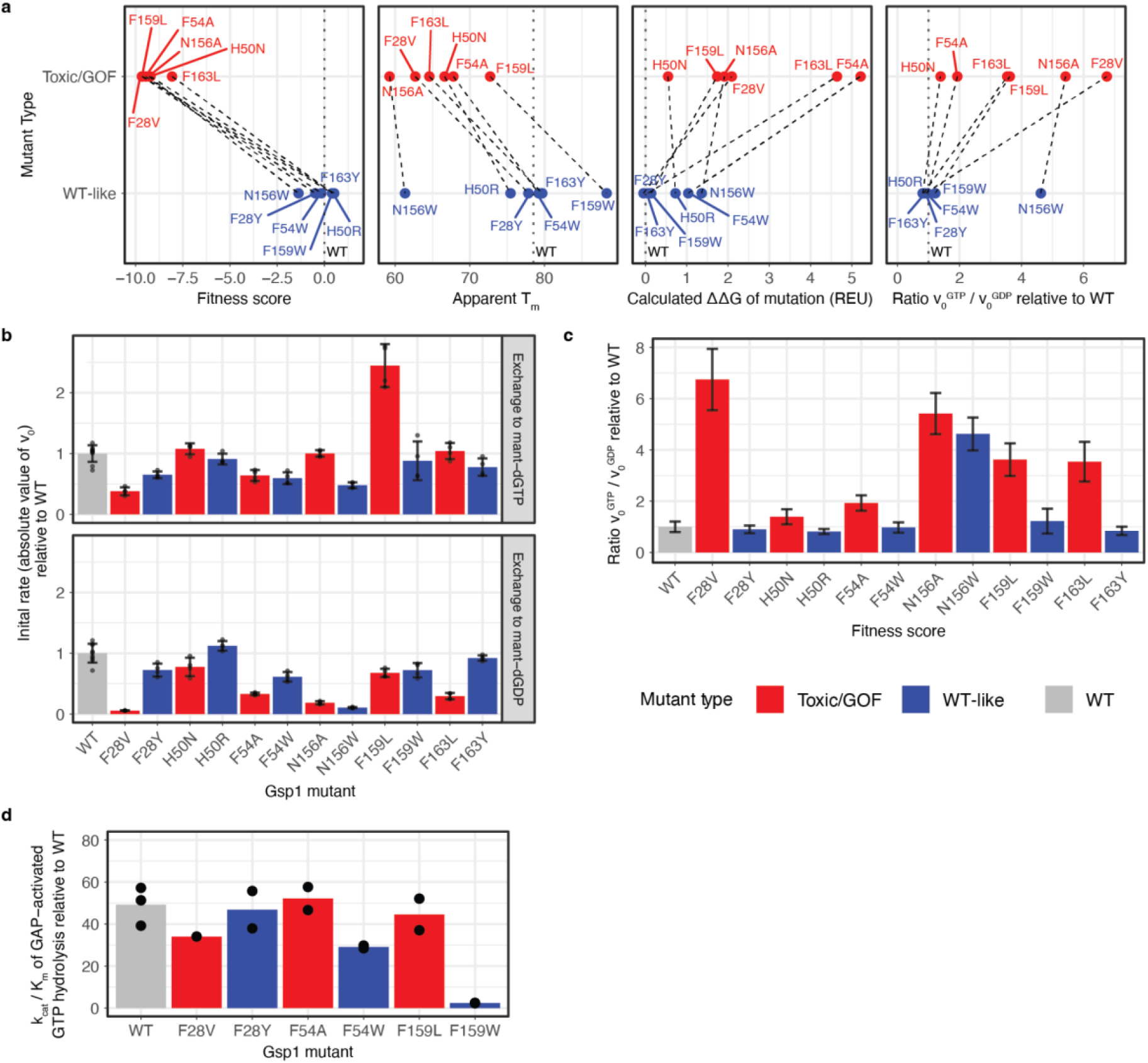
Biophysical properties of toxic/GOF and WT-like mutations. **a,** Comparison of toxic/GOF vs WT-like mutations for various parameters (from left to right): EMPIRIC fitness score, apparent *T_m_* as measured by irreversible CD melts, calculated ΔΔG for the GDP-bound structure (*H. sapiens* Ran-GDP, PDB ID: 3GJ0) using Rosetta, and the change in preference for GTP over GDP relative to WT as measured using the GEF-mediated nucleotide exchange assay. Variants colored by category: Toxic/GOF (red) and WT-like (blue). WT values shown as black dotted line. Colored dotted lines connect mutants at the same sequence position. **b,** Initial rates of GEF-mediated nucleotide exchange, normalized to the WT values. Measurements performed with n >= 4 replicates. Error bars represent the standard deviations of v0 measurements propagated across the division operator (Methods). **c,** Relative change in nucleotide preference, calculated as the ratio of initial rate of exchange to GTP divided by the initial rate of exchange to GDP, normalized to the wild-type ratio. Error bars represent the standard deviations of *v*_0_ measurements propagated across the division operator. **d,** Catalytic efficiency 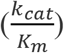 of GAP-activated GTP hydrolysis, normalized to the WT value. Individual replicates are shown as points on each bar.

**Figure S7.**
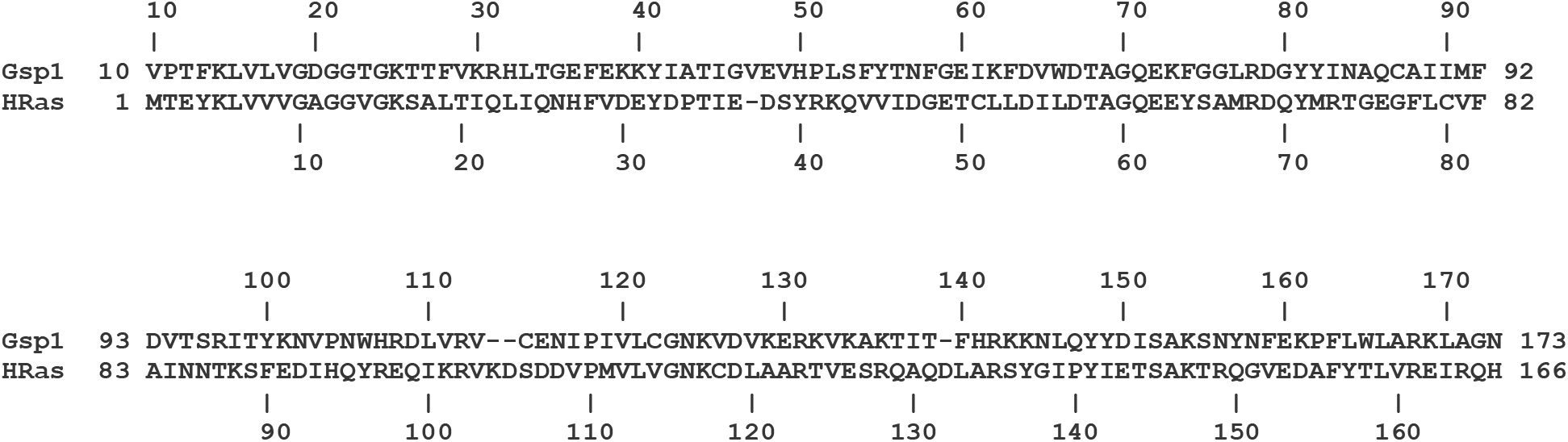
Sequence alignment of Gsp1 – H-Ras based on structural alignment.

**Figure S8.**
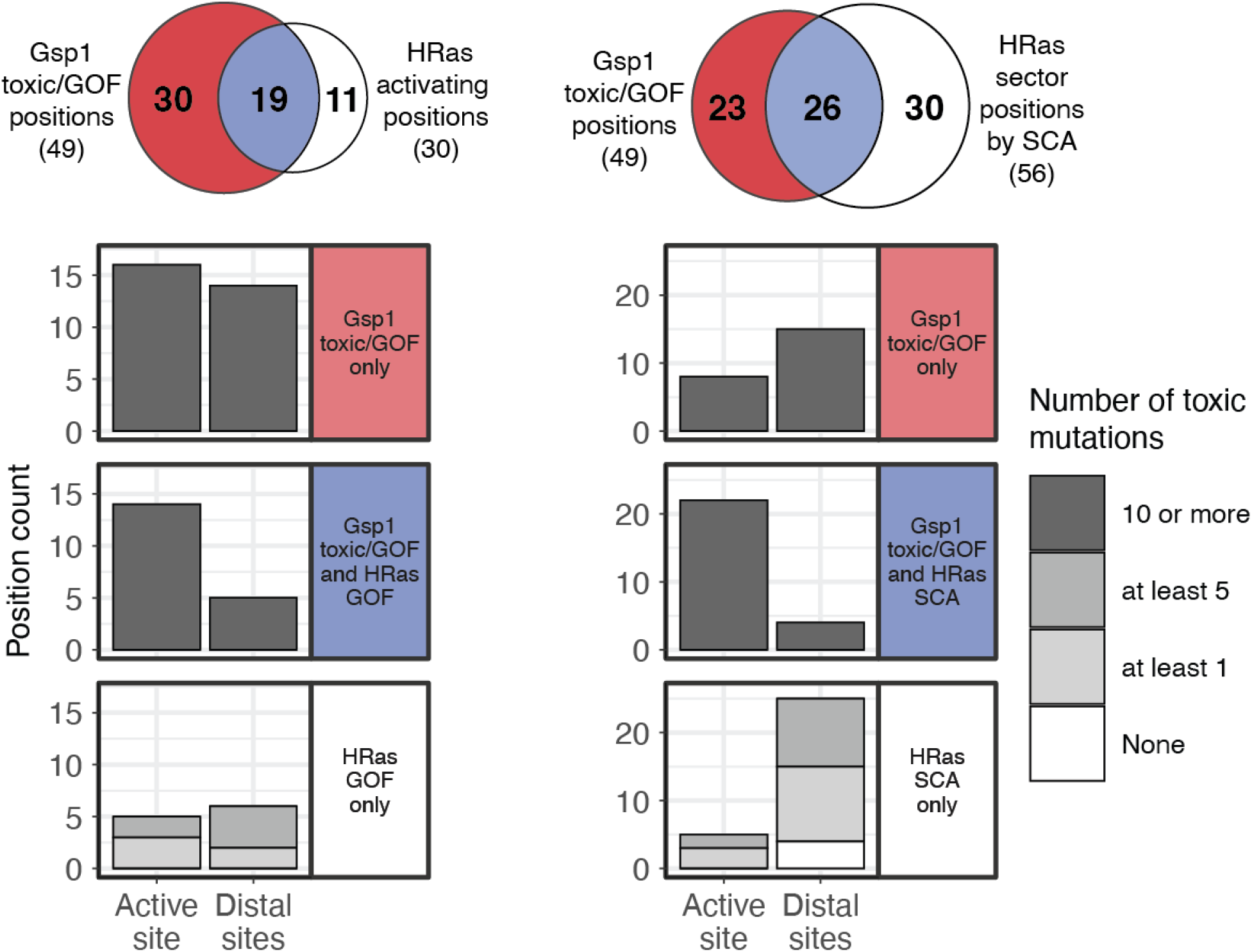
Locations of functional positions identified by the Gsp1 generalized EMPIRIC assay, HRas activation assay, or HRas statistical coupling analysis. Active site defined as in **Fig. 1**, including the canonical G1-5 regions conserved across Ras-superfamily GTPases, as well as residues included in the expanded definitions for Ran/Gsp1 based on structural analysis (**Fig. S1b** and Methods). Venn diagrams are as in **Fig. 4e**, **f** and repeated here for comparison. Bar graphs underneath indicate number of positions in each of the categories (red, blue, and white) from the Venn diagram. Bars are shaded by number of toxic/GOF mutations in the Gsp1 generalized EMPIRIC assay.

## References

1. Motlagh, H. N., Wrabl, J. O., Li, J. & Hilser, V. J. The ensemble nature of allostery. Nature 508, 331–339 (2014).

2. Faure, A. J. et al. Mapping the energetic and allosteric landscapes of protein binding domains. Nature 604, 175–183 (2022).

3. Piazza, I. et al. A Map of Protein-Metabolite Interactions Reveals Principles of Chemical Communication. Cell vol. 172 358–372.e23 (2018).

4. Hidalgo, F. et al. A saturation-mutagenesis analysis of the interplay between stability and activation in Ras. Elife 11, (2022).

5. Bandaru, P. et al. Deconstruction of the Ras switching cycle through saturation mutagenesis. Elife 6, 727 (2017).

6. Perica, T. et al. Systems-level effects of allosteric perturbations to a model molecular switch. Nature 1–6 (2021).

7. Jameson, N., Gavagan, M. & Zalatan, J. G. A kinetic mechanism for systems-level behavior in GTPase signaling. Trends Biochem. Sci. (2022) doi:10.1016/j.tibs.2022.01.006.

8. Dokholyan, N. V. Controlling Allosteric Networks in Proteins. Chem. Rev. 116, 6463–6487 (2016).

9. Kuriyan, J. & Eisenberg, D. The origin of protein interactions and allostery in colocalization. Nature 450, 983–990 (2007).

10. Kholodenko, B. N. Cell-signalling dynamics in time and space. Nat. Rev. Mol. Cell Biol. 7, 165–176 (2006).

11. Cherfils, J. & Zeghouf, M. Regulation of small GTPases by GEFs, GAPs, and GDIs. Physiol. Rev. 93, 269–309 (2013).

12. Vetter, I. R. & Wittinghofer, A. The guanine nucleotide-binding switch in three dimensions. Science 294, 1299–1304 (2001).

13. Ostrem, J. M., Peters, U., Sos, M. L., Wells, J. A. & Shokat, K. M. K-Ras(G12C) inhibitors allosterically control GTP affinity and effector interactions. Nature 503, 548–551 (2013).

14. Eisenberg, D., Marcotte, E. M., Xenarios, I. & Yeates, T. O. Protein function in the post-genomic era. Nature 405, 823–826 (2000).

15. Fowler, D. M. & Fields, S. Deep mutational scanning: a new style of protein science. Nat. Methods 11, 801–807 (2014).

16. Dunham, A. S. & Beltrao, P. Exploring amino acid functions in a deep mutational landscape. Mol. Syst. Biol. 17, e10305 (2021).

17. Stiffler, M. A., Hekstra, D. R. & Ranganathan, R. Evolvability as a function of purifying selection in TEM-1 β-lactamase. Cell 160, 882–892 (2015).

18. Roscoe, B. P., Thayer, K. M., Zeldovich, K. B., Fushman, D. & Bolon, D. N. A. Analyses of the effects of all ubiquitin point mutants on yeast growth rate. J. Mol. Biol. 425, 1363–1377 (2013).

19. Hietpas, R. T., Bank, C., Jensen, J. D. & Bolon, D. N. A. Shifting fitness landscapes in response to altered environments. Evolution 67, 3512–3522 (2013).

20. Hietpas, R., Roscoe, B., Jiang, L. & Bolon, D. N. A. Fitness analyses of all possible point mutations for regions of genes in yeast. Nat. Protoc. 7, 1382–1396 (2012).

21. Richards, S. A., Lounsbury, K. M. & Macara, I. G. The C Terminus of the Nuclear RAN/TC4 GTPase Stabilizes the GDP-bound State and Mediates Interactions with RCC1, RAN-GAP, and HTF9A/RANBP1*. J. Biol. Chem. 270, 14405–14411 (1995).

22. Zhou, J. et al. GEF-independent Ran activation shifts a fraction of the protein to the cytoplasm and promotes cell proliferation. Molecular Biomedicine 1, 18 (2020).

23. Holt, L. J. et al. Global analysis of Cdk1 substrate phosphorylation sites provides insights into evolution. Science 325, 1682–1686 (2009).

24. Neuwald, A. F., Kannan, N., Poleksic, A., Hata, N. & Liu, J. S. Ran’s C-terminal, basic patch, and nucleotide exchange mechanisms in light of a canonical structure for Rab, Rho, Ras, and Ran GTPases. Genome Res. 13, 673–692 (2003).

25. Weinert, B. T. et al. Lysine succinylation is a frequently occurring modification in prokaryotes and eukaryotes and extensively overlaps with acetylation. Cell Rep. 4, 842–851 (2013).

26. Swaney, D. L. et al. Global analysis of phosphorylation and ubiquitylation cross-talk in protein degradation. Nat. Methods 10, 676–682 (2013).

27. Rivoire, O., Reynolds, K. A. & Ranganathan, R. Evolution-Based Functional Decomposition of Proteins. PLoS Comput. Biol. 12, e1004817 (2016).

28. Goldbeter, A. & Koshland, D. E., Jr. An amplified sensitivity arising from covalent modification in biological systems. Proc. Natl. Acad. Sci. U. S. A. 78, 6840–6844 (1981).

29. Vetter, I. R. The Structure of the G Domain of the Ras Superfamily. in Ras Superfamily Small G Proteins: Biology and Mechanisms 1: General Features, Signaling (ed. Wittinghofer, A.) 25–50 (Springer Vienna, 2014).

30. Vetter, I. R., Arndt, A., Kutay, U., Görlich, D. & Wittinghofer, A. Structural view of the Ran-Importin beta interaction at 2.3 A resolution. Cell 97, 635–646 (1999).

31. Flynn, J. M. et al. Comprehensive fitness maps of Hsp90 show widespread environmental dependence. Elife 9, (2020).\

32. Gietz, R. D. & Schiestl, R. H. High-efficiency yeast transformation using the LiAc/SS carrier DNA/PEG method. Nat. Protoc. 2, 31–34 (2007).

33. Studier, F. W. Protein production by auto-induction in high density shaking cultures. Protein Expr. Purif. 41, 207–234 (2005).

34. Smith, S. J. M. & Rittinger, K. Preparation of GTPases for structural and biophysical analysis. Methods Mol. Biol. 189, 13–24 (2002).

35. Malaby, A. W. et al. Methods for analysis of size-exclusion chromatography–small-angle X-ray scattering and reconstruction of protein scattering. J. Appl. Crystallogr. 48, 1102–1113 (2015).

36. Joosten, R. P., Long, F., Murshudov, G. N. & Perrakis, A. The PDB_REDO server for macromolecular structure model optimization. IUCrJ 1, 213–220 (2014).

37. Touw, W. G. et al. A series of PDB-related databanks for everyday needs. Nucleic Acids Research vol. 43 D364–D368 (2015).

38. Chaudhury, S., Lyskov, S. & Gray, J. J. PyRosetta: a script-based interface for implementing molecular modeling algorithms using Rosetta. Bioinformatics 26, 689–691 (2010).

39. Levy, E. D. A simple definition of structural regions in proteins and its use in analyzing interface evolution. J. Mol. Biol. 403, 660–670 (2010).

40. Tien, M. Z., Meyer, A. G., Sydykova, D. K., Spielman, S. J. & Wilke, C. O. Maximum allowed solvent accessibilites of residues in proteins. PLoS One 8, e80635 (2013).

41. Vetter, I. R., Nowak, C., Nishimoto, T., Kuhlmann, J. & Wittinghofer, A. Structure of a Ran-binding domain complexed with Ran bound to a GTP analogue: implications for nuclear transport. Nature 398, 39–46 (1999).

42. Chook, Y. M. & Blobel, G. Structure of the nuclear transport complex karyopherin-beta2-Ran x GppNHp. Nature 399, 230–237 (1999).

43. Scheffzek, K., Klebe, C., Fritz-Wolf, K., Kabsch, W. & Wittinghofer, A. Crystal structure of the nuclear Ras-related protein Ran in its GDP-bound form. Nature 374, 378–381 (1995).

44. Renault, L., Kuhlmann, J., Henkel, A. & Wittinghofer, A. Structural basis for guanine nucleotide exchange on Ran by the regulator of chromosome condensation (RCC1). Cell 105, 245–255 (2001).

45. Park, H. et al. Simultaneous Optimization of Biomolecular Energy Functions on Features from Small Molecules and Macromolecules. J. Chem. Theory Comput. 12, 6201–6212 (2016).

46. Frenz, B. et al. Prediction of Protein Mutational Free Energy: Benchmark and Sampling Improvements Increase Classification Accuracy. Front Bioeng Biotechnol 8, 558247 (2020).

47. Ó Conchúir, S. et al. A Web Resource for Standardized Benchmark Datasets, Metrics, and Rosetta Protocols for Macromolecular Modeling and Design. PLoS One 10, e0130433 (2015).

48. Kellogg, E. H., Leaver-Fay, A. & Baker, D. Role of conformational sampling in computing mutation-induced changes in protein structure and stability. Proteins 79, 830–838 (2011).

49. Grant, B. J., Rodrigues, A. P. C., ElSawy, K. M., McCammon, J. A. & Caves, L. S. D. Bio3d: an R package for the comparative analysis of protein structures. Bioinformatics 22, 2695–2696 (2006).

50. Lockless, S. W. & Ranganathan, R. Evolutionarily conserved pathways of energetic connectivity in protein families. Science 286, 295–299 (1999).

