## Supplementary File 1 for "A complete allosteric map of a GTPase switch in its native network"

**Supplementary File Table of Contents**

Supplementary Table 1
Supplementary Figures 1 and 2
Titles and Legends of Supplementary Source Files

### Supplementary Table

**Supplementary Table 1. Overlap of positions annotated as toxic/GOF in Gsp1, activating in H-Ras, or part of an H-Ras sector by SCA**. Definitions for categories are from Gsp1 analysis, shown in **Fig. 4**. Sets of H-Ras sites for each category were constructed based on a structural alignment, and the subset of sites labeled as activating (Hidalgo et al. 2022 *eLife*) or as part of the SCA sector (Rivoire et al. 2016 *PLoS Computational Biology*) are listed.

| Gsp1 residue category (Fig. 4) | Toxic/GOF positions in Gsp1 (10 or more toxic mutations) | Activating positions in H-Ras (>=2 activating mutations) | Sector positions in H-Ras (from SCA analysis) |
|---|---|---|---|
| Active site regions | G19, D20, G21, G22, T23, G24, K25, T26, G35, E36, F37, A43, T44, I45, G46, D67, T68, A69, G70, Q71, E72, G75, L77, R78, G80, Y81, K125, D127, S152, A153 | G12, G13, V14, K16, A18, P34, T58, A59, G60, Q61, E63, R68, N116, K117, D119, L120, S145, A146, K147 | G10, A11, V14, G15, K16, S17, F28, Y32, P34, T35, I36, D57, T58, A59, G60, Q61, E62, E63, Y64, R68, Y71, N116, K117, D119, S145, A146, K147 |
| Distal sites affecting switching | F28, H32, G35, H50, F54, N156, Y157, F159, F163 | L19, Q22, L23, V152, F156 | Q22, L23, A134, F156 |
| PTM sites in Gsp1 | K25, K101, K125, S155 | K16, K117 | K16, K117 |
| Regulator Interface | G21, G22, G35, E36, F37, A43, I45, T56, G59, A69, Q71, E72, G75, D93, S96, R97, T99, K101, K132, V133, N156, F159, E160 | G12, G13, P34, A59, Q61, E63, A83, V152 | F28, Y32, P34, I36, A59, Q61, E62, E63, Y64, A83, Q99, R123, V125 |

### Supplementary Figures

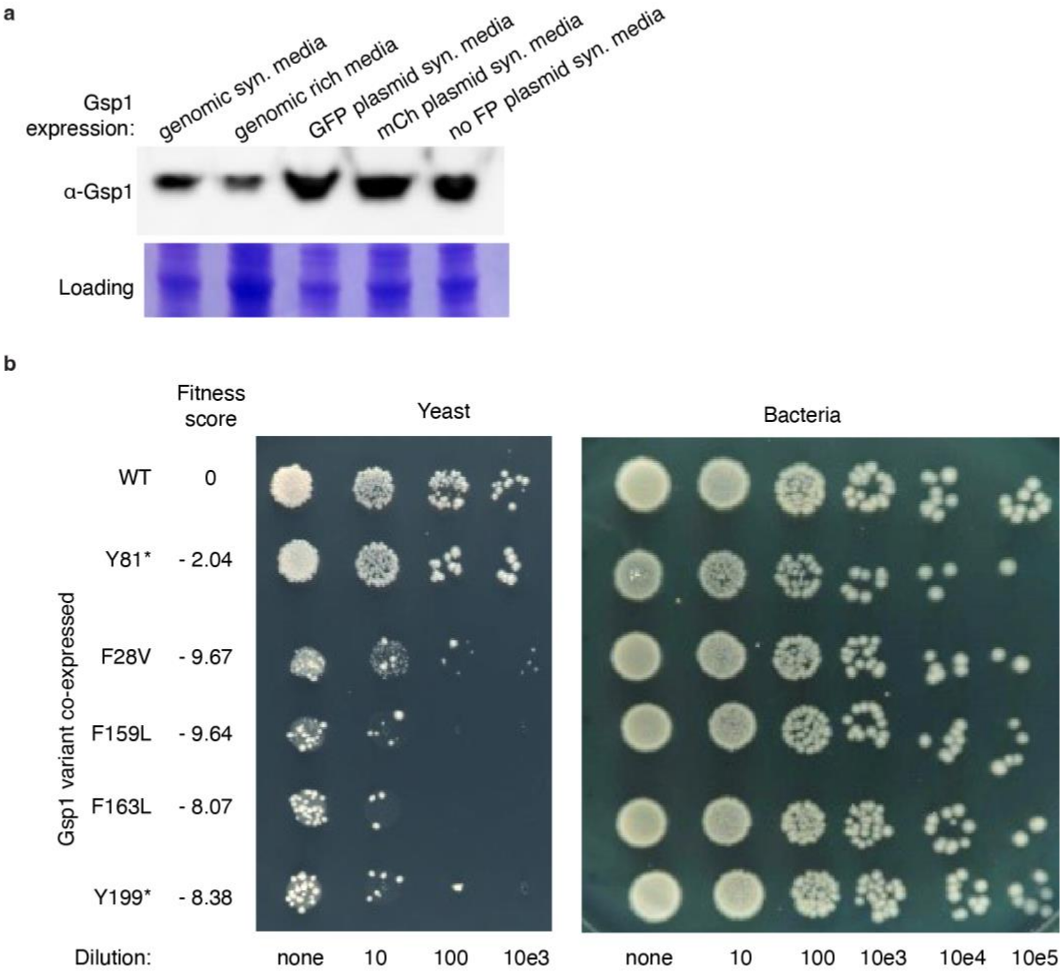


**Supplementary Figure 1. Verification of EMPIRIC plasmid expression and growth phenotypes of individual Gsp1 variants. a,** Western blot comparing the expression of genomic Gsp1 (lanes 1 and 2) to that of Gsp1 expressed from an EMPIRIC plasmid driven by its native promoter (lanes 3-5) in the presence of a GFP marker (lane 3), mCherry marker (lane 4) or no marker (lane 5). **b,** Dilution series for individual Gsp1 variants expressed from an EMPIRIC plasmid. *S. cerevisiae* (left) and *E. coli* (right) cells were transformed with equal amounts of plasmid and subsequently spotted onto selective plates in a dilution series. Series are shown for WT, one internal STOP codon mutant (Y81*), three toxic/GOF substitution variants (F28V, F159L, F163L), and one C-terminal extension STOP codon mutant (Y199*). Corresponding fitness scores from the EMPIRIC assay are provided. Bacterial dilutions show that the EMPIRIC plasmid itself is not generally toxic.

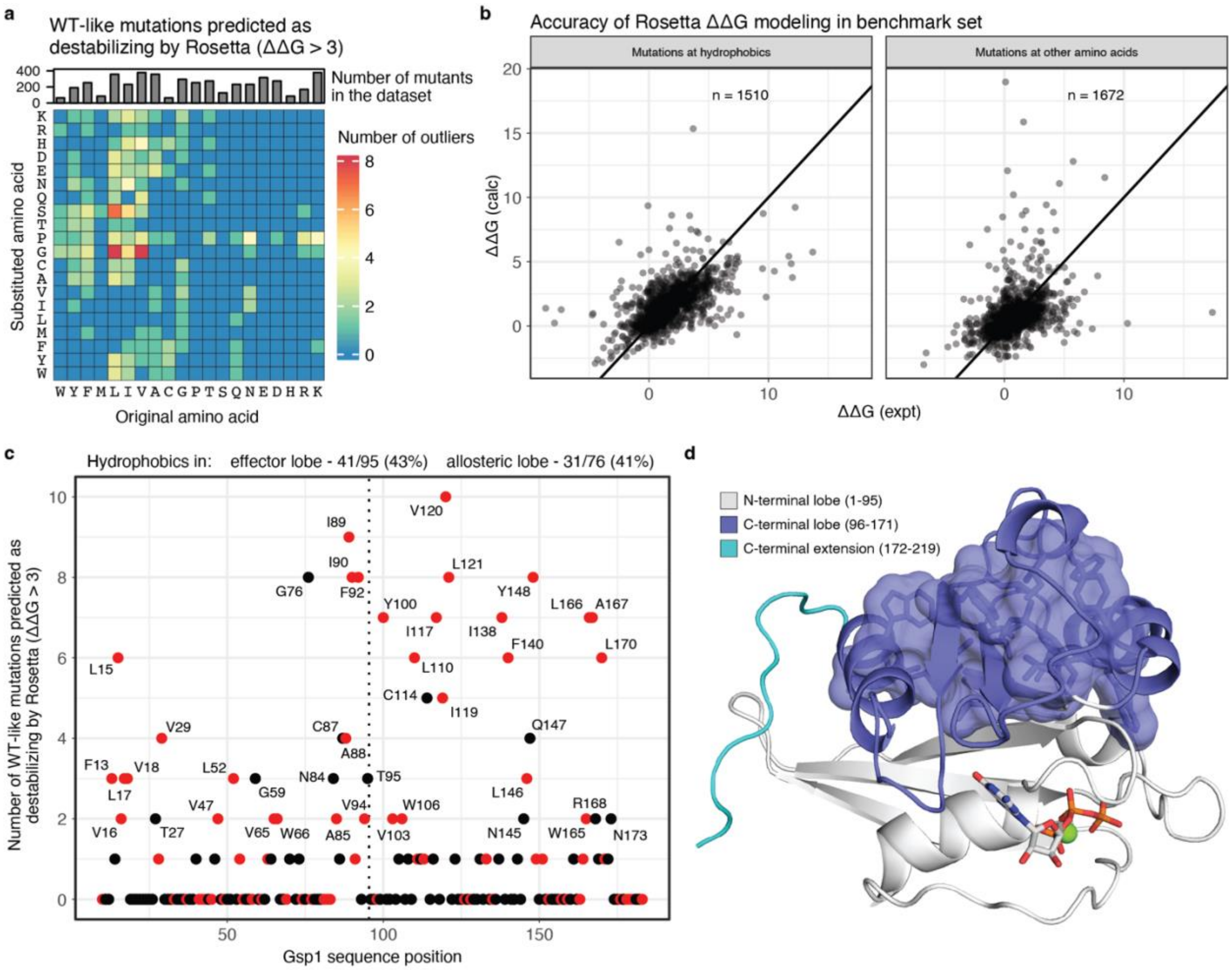


**Supplementary Figure 2. WT-like mutations predicted as destabilizing are predominantly at hydrophobic positions in the C-terminal lobe. a,** Heatmap showing the number of mutations with WT-like fitness scores that are predicted as destabilizing (ΔΔG > 3 Rosetta Energy Units, REU) for Gsp1-GTP, broken down by original and substituted amino acid. **b,** Experimentally measured and computed ΔΔG values in a large benchmark dataset separated by mutations at positions with a hydrophobic WT amino acid residue (W, Y, F, M, L, I, V, A), vs. those at all other residues. **c,** Number of mis-predicted WT-like mutations as in panel (a), ordered by Gsp1 sequence position. Positions with two or more mutations mis-predicted are labeled. Mutations at hydrophobic positions shown in red. A large set of mutations at core hydrophobic residues in the C-terminal lobe of Gsp1 are predicted as destabilizing but have WT-like fitness scores, a discrepancy that cannot be explained by poor method accuracy of Rosetta given high performance on similar mutations in a large benchmark dataset as seen in panel (c). **d,** Structural representation of Gsp1-GTP (PDB ID: 3M1I) showing the N-terminal lobe (white), C-terminal lobe (purple), and a stretch of C-terminal extension residues (cyan). Hydrophobic side chains comprising the C-terminal lobe core (red dots after sequence position 87 in panel (d)) are shown in sticks and surface representation. Nucleotide shown in white sticks and $Mg^{2+}$ cofactor shown as a green sphere.

**Titles and Legends of Supplementary Source Files**

**Source File 1 Gsp1 fitness scores with bins and raw read counts**

Column definitions:

- *mutant*: descriptor of the allele (e.g. F28V)
- *aa_from*: wild-type amino acid (e.g. F)
- *position*: sequence position of the mutation (e.g. 28)
- *aa_to*: substituted amino acid (e.g. V)
- *counts_0gen*: number of reads corresponding to the allele at the beginning of selection
- *counts_6gen*: number of reads corresponding to the allele after 6 generations of selection
- *score*: computed fitness score, a log2-transformed changes in variant abundance relative to wild-type
- *bin*: assigned score bin
- *low_reads_flag*: boolean flag stating whether the allele had low read counts in the initial sample (< 2% of the average allele's number of reads)

**Source File 2 Gsp1 ΔΔG data**

Column definitions:

- *mutation*: descriptor of the mutation (e.g. F28V)
- *aa_from*: wild-type amino acid (e.g. F)
- *aa_to*: substituted amino acid (e.g. V)
- *pdb_id*: PDB ID of the crystal structure used for the calculation
- *species*: organism of the gene used for determination of the crystal structure
- *pos_Sc*: sequence position number of the corresponding residue in *S. cerevisiae* Gsp1
- *aa_Sc*: wild-type amino acid of the corresponding residue in *S. cerevisiae* Gsp1
- *pos_Hs*: sequence position number of the corresponding residue in *H. sapiens* Ran
- *aa_Sc*: wild-type amino acid of the corresponding residue in *H. sapiens* Ran
- *ddg*: computed ΔΔG of the mutation, in Rosetta Energy Units (REU), scaled so that 1 REU ~ 1 kcal/mol based on the benchmark data

**Source File 3 Benchmark ΔΔG data**

Column definitions:

- *record_id*: unique ID used in this study for the benchmark mutation
- *pdb*: PDB ID of the crystal structure used for the calculation
- *mutation_full*: long form ID for the mutation made (e.g. A T 100 G corresponds to mutating the Threonine at position 100 in chain A to Glycine)
- *chain*: one-letter ID for the crystal structure chain used for the calculation
- *mutation*: short form descriptor of the mutation (e.g. T100G)
- *position*: sequence position of the mutation (e.g. 100)
- *ddg_expt*: experimental ΔΔG of the mutation, in kcal/mol
- *ddg_calc*: computed ΔΔG of the mutation, in unscaled Rosetta Energy Units (REU)
- *ddg_calc_adj*: computed ΔΔG of the mutation, in REU, scaled down so that 1 REU ~ 1 kcal/mol
